# Therapeutically actionable PAK4 is amplified, overexpressed and involved in bladder cancer progression

**DOI:** 10.1101/740316

**Authors:** Darshan S. Chandrashekar, Balabhadrapatruni V. S. K. Chakravarthi, Alyncia D. Robinson, Joshua C. Anderson, Sumit Agarwal, Sai Akshaya Hodigere Balasubramanya, Marie-Lisa Eich, Akhilesh Kumar Bajpai, Sravanthi Davuluri, Maya S. Guru, Arjun S. Guru, Gurudatta Naik, Deborah L. Della Manna, Kshitish K. Acharya, Shannon Carskadon, Upender Manne, David K. Crossman, James E. Ferguson, William E. Grizzle, Nallasivam Palanisamy, Christopher D. Willey, Michael R. Crowley, George J Netto, Eddy S. Yang, Sooryanarayana Varambally, Guru Sonpavde

## Abstract

Muscle-invasive bladder carcinomas (MIBCs) are aggressive genitourinary malignancies. Disease incidence and survival rates vary based on aggressiveness and treatment options. Metastatic urothelial carcinoma of the bladder is generally incurable by current chemotherapy and leads to early mortality. For a minority (∼20%) of patients, T-cell checkpoint inhibitors provide durable benefits following prior platinum therapy. Recent studies have identified molecular subtypes of MIBCs with different sensitivities to frontline therapy, suggesting heterogeneity in these tumors and pointing to the importance of molecular characterization of MIBCs to provide effective treatment. We have performed multi-omic profiling of the kinome to identify therapeutic targets that are overexpressed in a subset of BLCAs. Our analyses revealed amplification and overexpression of P21 (RAC1) activated kinase 4 (PAK4) in a subset of BLCAs. For these tumors, multiplex kinase assay profiling identified corresponding PAK4 target substrates. By performing experiments using cultured bladder cancer cells, we confirmed the role of PAK4 in BLCA cell proliferation and invasion. Furthermore, our studies showed that a PAK4 inhibitor was effective in curtailing growth of BLCA cells. Transcriptomic analyses identified elevated expression of another kinase, Protein Tyrosine Kinase 6 (PTK6), upon treatment with a PAK4 inhibitor. Similarly, RNA interference of PAK4 led to elevated expression of PTK6. Treatment with a combination of kinase inhibitors (vandetanib and dasatinib) showed enhanced sensitivity compared to either drug alone. Thus, PAK4 may be therapeutically actionable for a subset of MIBC patients with amplified and/or overexpressed PAK4 in their tumors. Our results also indicate that combined inhibition of PAK4 and PTK6 may overcome resistance to PAK4. These observations warrant clinical investigations with selected BLCA patients.

## Introduction

Bladder cancer (BLCA) is the ninth-most common malignancy worldwide (1). Most BLCAs are urothelial carcinomas, and metastatic urothelial carcinoma is generally incurable by current cisplatin-based first-line chemotherapy, leading to early mortality (2). Perioperative cisplatin-based combination chemotherapy for localized muscle-invasive BLCA (MIBC) modestly improves survival; however, approximately half of all patients progress rapidly with distant metastatic disease (3, 4). For a minority (∼20%) of patients with prior platinum therapy, T-cell checkpoint inhibitors have recently provided durable benefits as first-line therapy for cisplatin-ineligible patients with high tumor PD-L1 protein expression (5-10). Ongoing clinical trials are evaluating the combination of PD1/PD-L1 inhibitors with platinum-based first-line chemotherapy or with CTLA-4 inhibitors. Given the molecular heterogeneity of this disease, rational therapeutic targeting guided by somatic genomic alterations may hold promise.

Various molecular alterations are involved in the progression of aggressive BLCAs. Recent studies have identified, for MIBCs, molecular subtypes based on gene expression that have different sensitivities to frontline therapy, suggesting heterogeneity in these tumors and the importance of molecular characterization of the cancers to provide effective treatment (11-14). Dysregulation of the transcriptional maintenance system is believed to be a cause for malignancy (15, 16). The advent of new technologies allows molecular analyses of BLCAs and enhances the promise of targeted therapy and personalized medicine (17, 18).

Kinases, which are common drivers of malignancies, are potentially actionable for therapy. Kinase inhibitors, which are readily manufactured, are approved to treat various malignancies. Inhibitors of fibroblast growth factor receptor (FGFR), ERBB2, and mTOR kinase have activity against tumors harboring genomic alterations in their respective genes (19-21). However, the primary kinase drivers of growth of urothelial carcinomas are unclear. Thus, we performed genomic, transcriptomic, and kinomic analyses of MIBC to identify aberrations in the kinome.

## Results

### Multi-platform kinase analysis of tumor and normal samples identifies PAK4 as the primary amplified and overexpressed kinase gene

Targeted kinome gene sequencing of 24 MIBC samples and their matched normal tissues led to identification of somatic mutations/indels and somatic copy number alterations. Overall, 24 kinases harbored somatic mutations/indels in two or more MIBC samples. TTN, OBSCN, EPHA5, FASTK, and MAST1 were most frequently mutated. For these samples, copy number analyses found PAK6, TTN, NEK1, and CDK17 to be predominantly deleted (≥4 of 24); RIPK3, PAK4, and TTBK2 were predominantly amplified (≥2 of 24) (**Figure 1A**,**B**). Fluorescence in situ hybridization (FISH) using a PAK4 locus-specific probe revealed copy number gains in BLCA cells (**Figure 1C**). Kinase gene expression estimated by NanoString assays was analyzed for all frequently mutated/altered genes found by kinome sequencing analysis. Among the amplified genes, PAK4 [Fold change: 1.75, P-value: 0.0025] showed up-regulation in MIBC samples compared to matched normal tissues (**Figure 1D**); among deleted genes, TTN [fold change: −5.96, P-value: 1E-8], CDK17 [fold change: −1.95, P-value: 1.1E-7], and NEK1 [fold change: −2.26, P-value: 3.2E-5] showed down-regulation.

**Figure 1:**
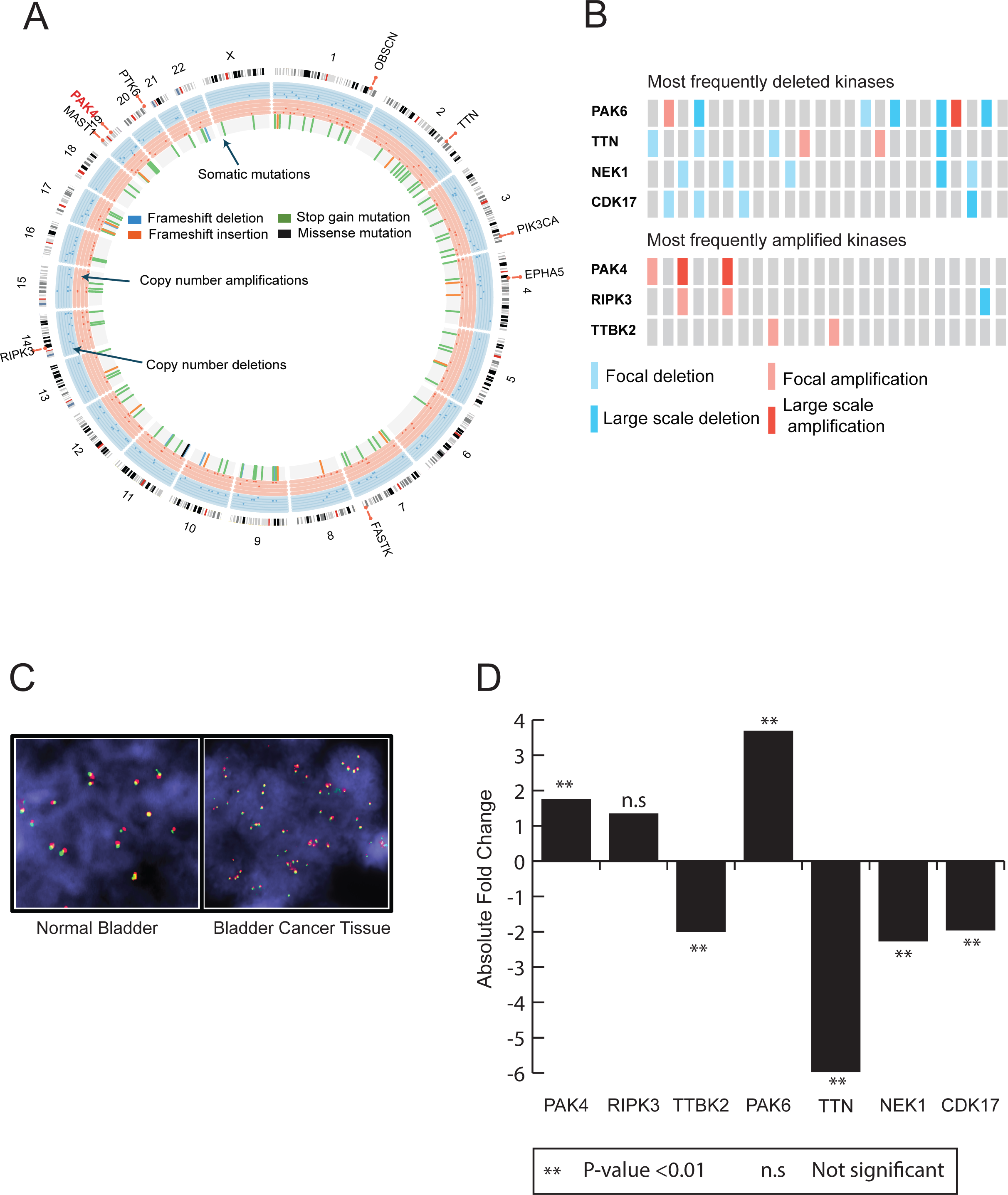
Multi-platform kinome analysis of 24 MIBCs and their matched normal tissues. (A) Circos plot showing kinome-wide somatic mutations, copy number alterations, and RNA-level expression. The outermost track is an ideogram running from chromosome 1 to chromosome X in a clockwise direction. The innermost track shows somatic mutations identified by both Mutect2 and Varscan2. The next two innermost tracks give an account of copy number amplifications and deletions identified by Varscan2. The subsequent two tracks summarize the RNA-level expression profile of kinases from a NanoString assay, in terms of proportion of samples with kinase over-/under-expression. (B) Copy number analysis identified the most frequently amplified (PAK4, RIPK3 and TTBK2) and deleted (PAK6, TTN, NEK1 and CDK17) genes. (C) FISH analysis of BLCA cells using a PAK4 locus-specific probe on chromosome 10q. Analysis showing multiple copies of PAK4 in BLCA cells (right); only two copies of PAK4 were present in normal bladder cells (left). (D) RNA expression levels of PAK4, RIPK3, TTBK2, PAK6, TTN, NEK1 and CDK17 in MIBCs (n=24) compared to matched normal tissues (n=20) using a customized NanoString platform.

### TCGA validation confirms PAK4 as an alteration in MIBC associated with higher stage and worse outcomes

We validated the most frequently amplified/deleted kinases (PAK6, TTN, NEK1, CDK17, RIPK3, PAK4, and TTBK2) using TCGA BLCA dataset via cBioPortal (**Supplementary Figure 1A**,**B**). PAK4 emerged as a biologically plausible driver gene, with 5% of TCGA BLCAs showing copy number amplification. Most PAK4-amplified TCGA BLCA samples (13 of 20) were of the luminal molecular subtype (luminal/luminal_papillary/luminal_infiltrated) (**Supplementary Table 1**). We also observed elevated expression of PAK4 in distinct histological, molecular subtypes and in advanced stages of the samples (**Figure 2A, Supplementary Figure 1C-E**). This was confirmed by qRT-PCR using MIBC RNA and normal bladder tissue RNA (n=11) (**Figure 2B**). There was also poorer survival (Log Rank P-value: 0.0473) of patients with PAK4 alterations [amplification/mRNA dysregulation] in BLCAs (**Supplementary Figure 2A**). Immunoblot analyses using PAK4 antibody showed that PAK4 protein expression was generally higher in BLCAs compared to normal bladder tissue **(Figure 2C**). We highlighted the PAK4-specific band, which was identified by the molecular weight and siRNA knockdown in bladder cells and by immunoblotting. Since the antibody produces multiple bands, we confirmed the specific band with PAK4 knockdown in VM-CUB1 cells (**Figure 2C, right panels**). We also measured PAK4 RNA and protein expression using various BLCA cell lines (**Figure 2D**,**E**).

**Figure 2:**
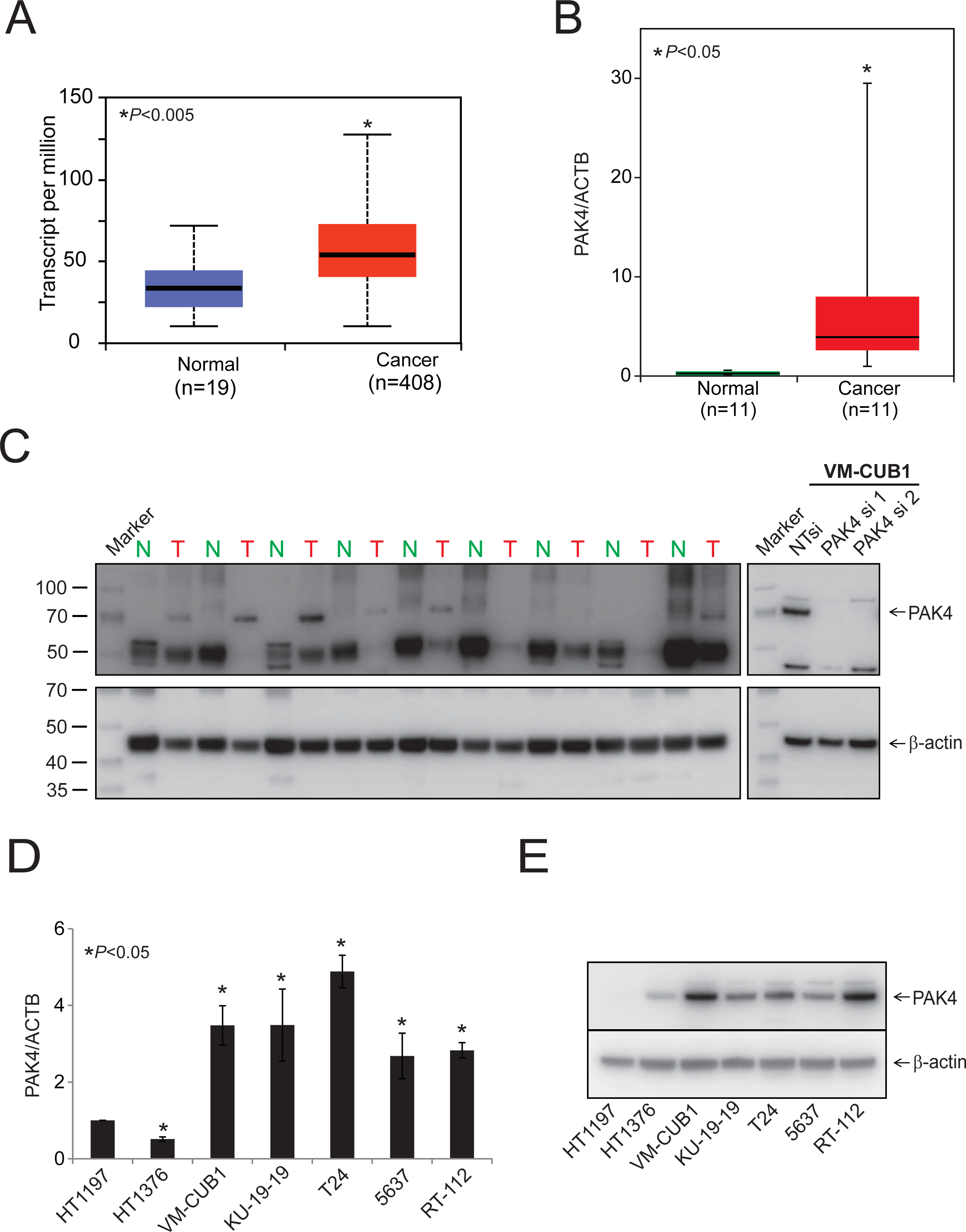
Elevated expression of PAK4 in aggressive bladder adenocarcinomas. (A) PAK4 gene expression from next-generation RNA sequencing data for normal and BLCA tissues (TCGA datasets). (B) qRT-PCR of PAK4 using RNA from matched normal and BLCA tissues. (C) Immunoblot analysis of PAK4 protein expression in matched normal and primary MIBC samples and PAK4 siRNA-treated VM-CUB1 cells using a PAK4 antibody and β-actin as a loading control. (D, E) qRT-PCR and Western blot analysis of PAK4 across BLCA cell lines. β-Actin was used as a loading control.

### Kinase assays show increased activity in BLCAs overexpressing PAK4

As kinases are frequently regulated post-transcriptionallyand their activity plays a key role, we measured kinase activity in tumor samples by an enzymatic kinomic assay and found phosphorylation changes for the four identified PAK4 substrates (**Figure 3A**), which were altered inter-patient between paired normal-tumor tissues (**Figure 3B**). Of the 24 patient tumors, 8 (33%) had high PKAS values (>2.0) (**Figure 3C**). For copy number variations (CNVs), 3 (37.5%) had amplifications, with two large-scale and one focal amplification.

**Figure 3:**
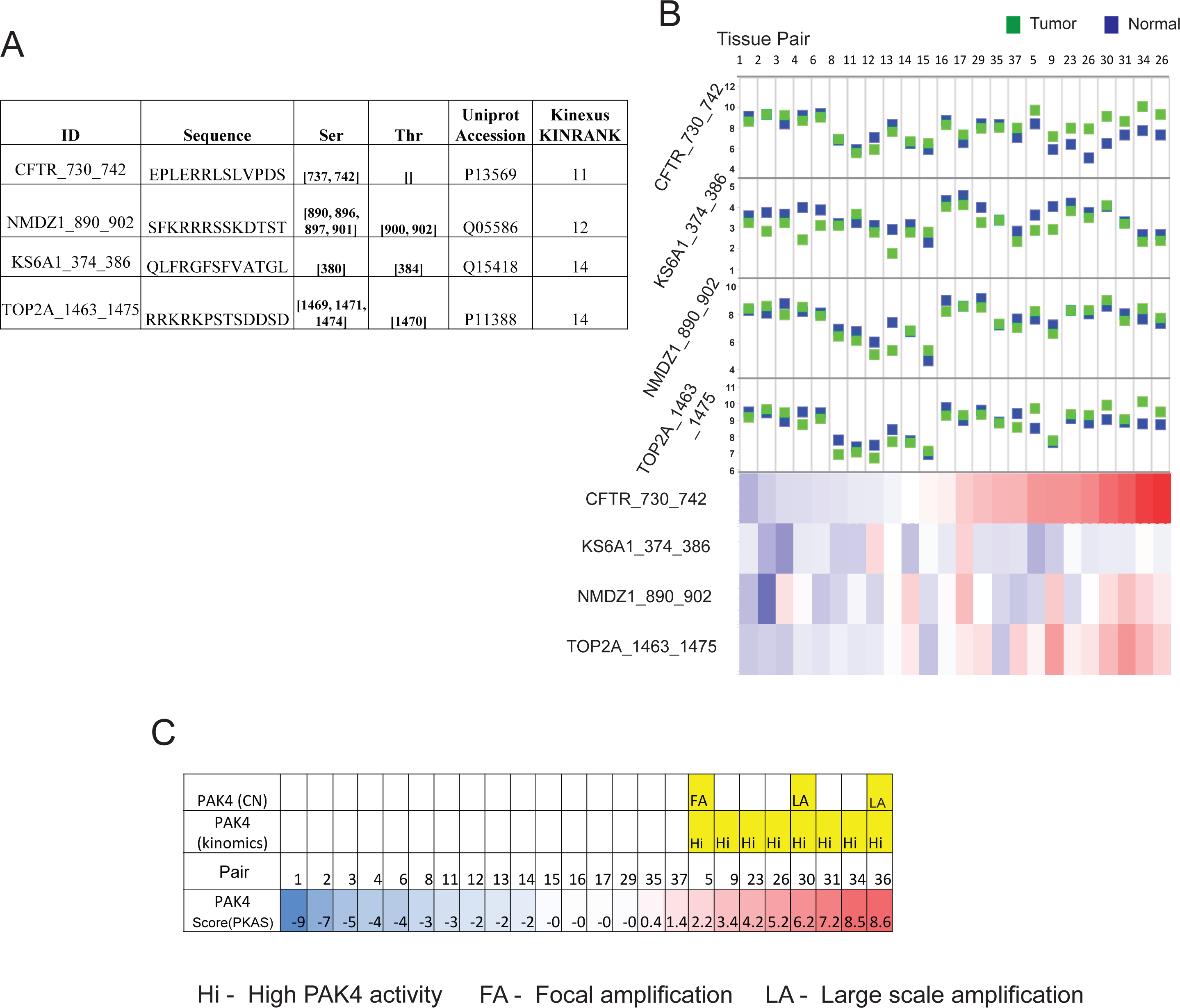
Elevated PAK4 activity in a subset of primary BLCAs. (A) The table displays peptides selected as PAK4 substrates and includes peptide sequence information and Kinexus scoring rank. (B) Overlays of peptide phosphorylation (end-level) for paired normal (blue) and tumor (green) activity of selected PAK4 substrates, with raw values plotted above and subtracted ‘tumor change’ values heatmapped below. (C) Rank orders by PKAS (bottom row) and highlighted PAK4 findings from other methods (yellow).

### PAK4 is involved in bladder cancer cell growth, invasion, and colony formation

Next, to investigate the role of PAK4 kinase in BLCA biology, we performed transient PAK4 knockdown in VM-CUB1 and RT-112 cells using two independent and specific siRNAs. To confirm the knockdown efficiency, we performed immunoblot analysis using protein lysates prepared after 72 hours of transient transfection (**Figure 4A-B, inset**). We evaluated cell proliferation, invasion, and colony formation using control and PAK4-knockdown cells. The cell proliferation assay using PAK4 knockdown cells indicated lower cell numbers (**Figures 4A-B**). VM-CUB1 cells with PAK4 knockdown showed less invasive potential in Boyden chamber Matrigel invasion assays (**Figure 4C**). (Note: One of the siRNAs did not show any phenotypic changesin RT-112 cells). Additionally, cells with PAK4 knockdown showed less colony formation (**Figure 4D**). These observations indicate an essential role for PAK4 in proliferation, invasion, and colony formation of BLCA cells.

**Figure 4:**
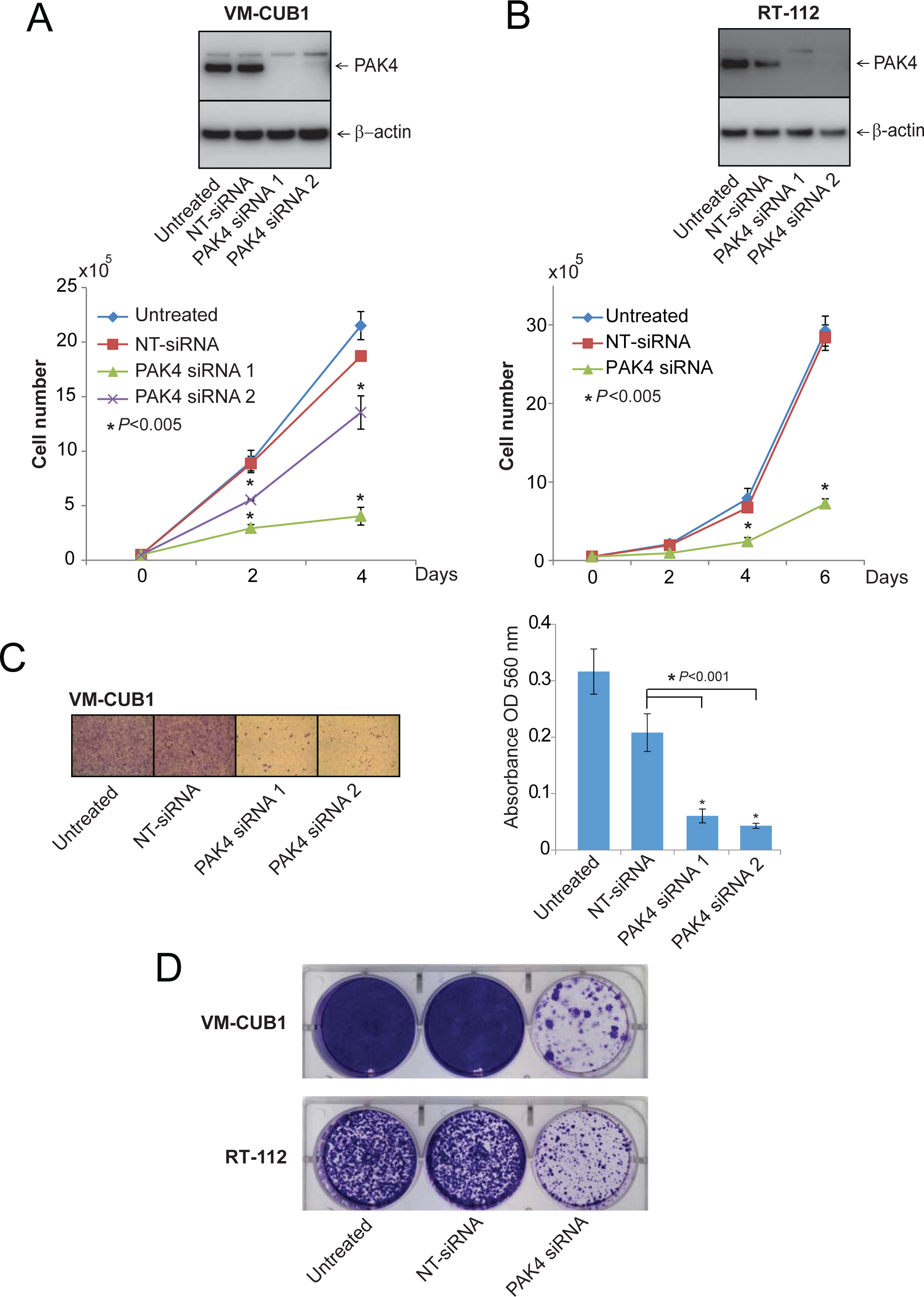
PAK4 involvement in BLCA cell proliferation and invasion. (A, B) Western blot analysis of BLCA cells with PAK4 transiently knocked down by treatment with either of two specific PAK4 siRNA duplexes. β-Actin was used as a loading control. Proliferation assay of these cells transfected with either of two PAK4 siRNA duplexes or NT siRNA. (C) Knockdown of PAK4 reduced VM-CUB1 cell invasion. Boyden chamber Matrigel invasion assays were performed using VM-CUB1 cells in which PAK4 was transiently knocked down using either of two specific siRNA duplexes. Untreated and NT siRNA-treated cells served as controls. Invaded cells were stained with crystal violet, and the absorbance was measured at 560 nm. (D) In a colony formation assay, the colony formation efficiency was reduced in PAK4 siRNA-treated cells as compared with untreated and NT siRNA controls.

### MicroRNAs miR-122 and miR-193 are involved in regulation of PAK4 expression

In addition to PAK4 amplification in 12% of BLCAs, there was upregulation of PAK4 in 50% of MIBCs. Hence, to test other forms of regulation, we investigated microRNAs that might target PAK4. Our analyses using Targetscan (22) suggested that microRNA-122 and 193 had binding sites at the 3’UTR of PAK4 (**Supplementary Figure 3A**). In various cancers, microRNA-122 and 193 are downregulated through epigenetic mechanisms (23-26). When these microRNAs were introduced into VM-CUB1 and RT-112 cells, there was downregulation of PAK4 expression (**Supplementary Figure 3B**); this was not caused by control microRNAs. Furthermore, addition of the microRNAs reduced colony formation (**Supplementary Figure 3C**). Thus, in a subset of bladder cancers, downregulation of PAK4 regulating microRNAs could lead to PAK4 overexpression.

### The PAK4 inhibitor reduces bladder cancer cell proliferation and colony formation

To evaluate the effect of PAK4 inhibition by a small molecule targeting PAK4, we performed proliferation and colony formation assays with VM-CUB1 and RT-112 cells that were untreated, treated with vehicle control, or treated with the PAK4 inhibitor (vandetanib) at various concentrations for 6 days. In the presence of the inhibitor, there was less cell proliferation (**Figure 5A-B**). Further, PAK4 at nanomolar concentrations reduced colony formation (**Figure 5C**). (Note that the degree of specificity of vandetanib for PAK4 could vary, as is the case for other kinase inhibitors). These results show that, for BLCAs, inhibition of PAK4 could be effective in reducing or blocking cancer growth.

**Figure 5:**
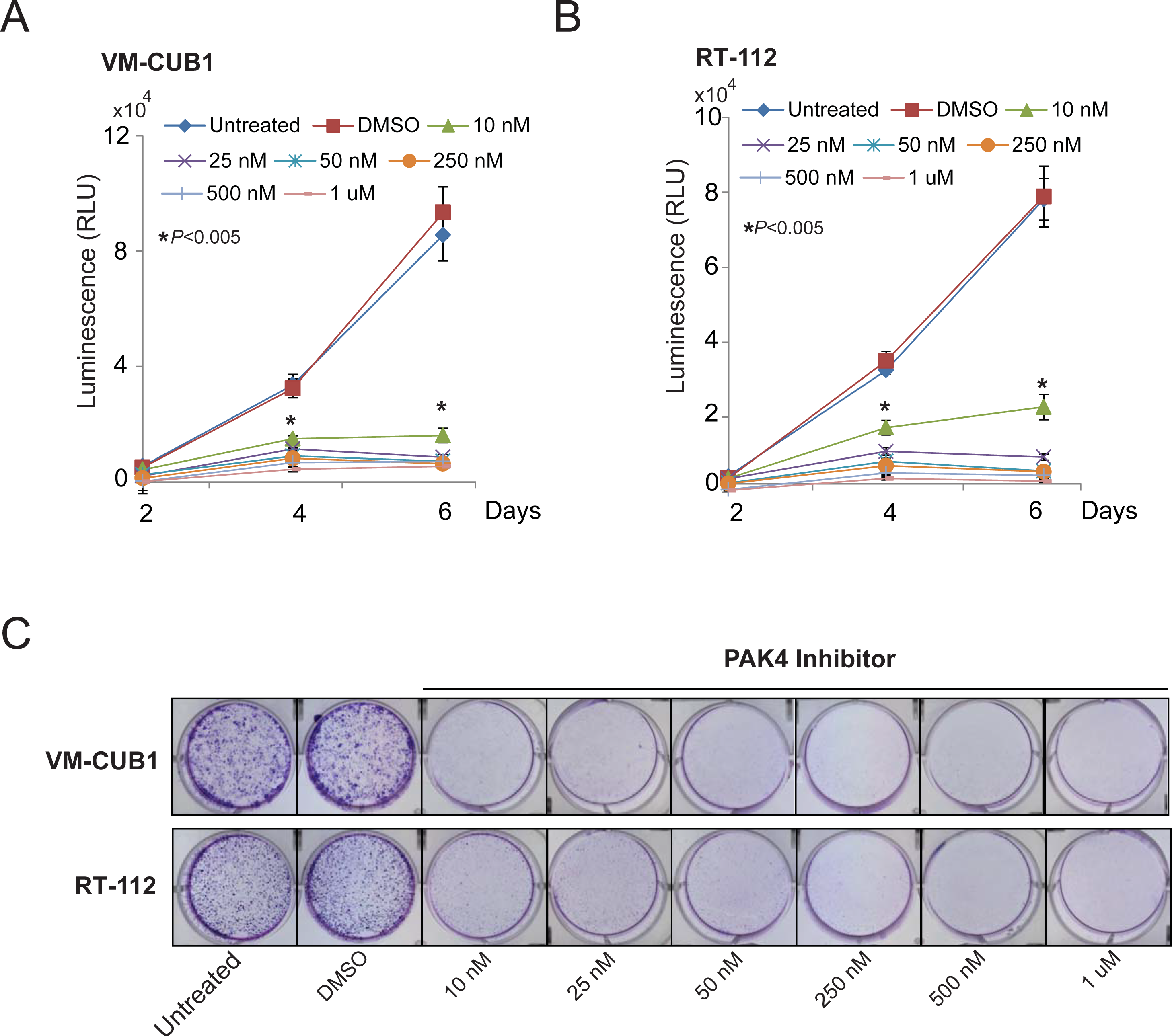
Inhibition of PAK4 by a small molecule inhibitor (PF-3758309) reduces BLCA cell proliferation. (A, B) cell proliferation and (C) colony formation assay images for VM-CUB1 and RT-112 cells treated with various concentrations of the PAK4 inhibitor.

### The global effects of PAK4 inhibition and PAK4 knockdown on bladder cancercells

To identify downstream targets of PAK4, whole transcriptome sequencing (RNA-seq) of VM-CUB1 cells treated with the PAK4 inhibitor or PAK4 siRNA was performed. Comparison of gene expression profiles for inhibitor-treated cells with those of DMSO-treated cells identified 166 up-regulated (123 protein-coding) and 259 down-regulated (196 protein-coding) genes. Similarly, 222 up-regulated (186 protein-coding) and 173 down-regulated (164 protein-coding) genes were identified from differential expression analysis of cells treated with PAK4 siRNA relative to NT siRNA (**Supplementary Table 2**). Those with absolute fold changes of ≥1.5 and P-values <0.05 were selected as differentially expressed genes [DEG] (**Figure 6 A**). Comparison of protein-coding DEGs from PAK4-inhibitor treated cells and PAK4 siRNA-treated cells showed 10 commonly up-regulated and 26 commonly down-regulated genes (**Figure 6B**).

**Figure 6:**
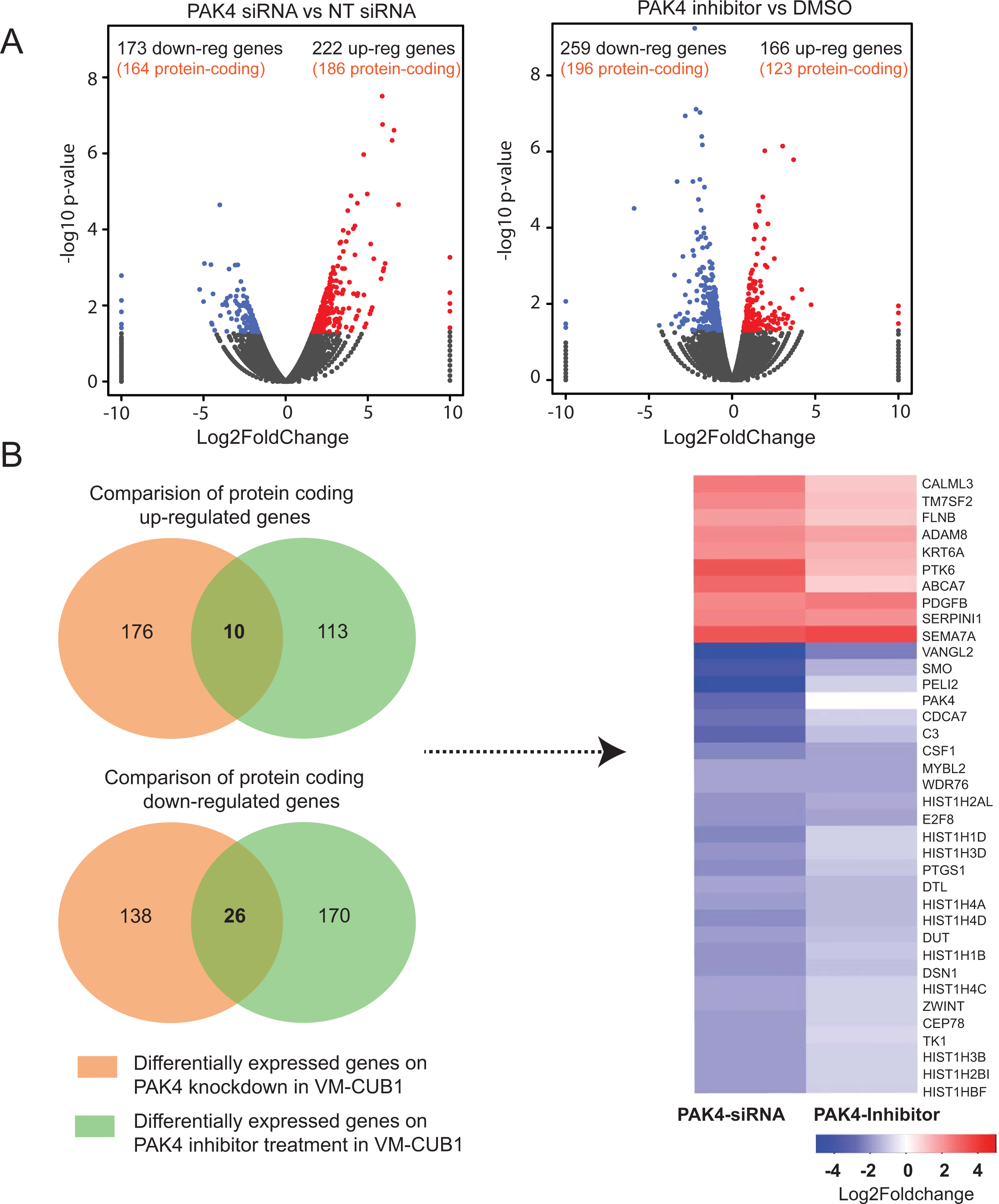
RNA-seq analysis of VM-CUB1 cells treated with PAK4 siRNA or PF-3758309. (A) Volcano plots showing the effect of PAK4 knockdown and PAK4 inhibition on global gene expression in VM-CUB1 cells. Up-regulated genes are shown in red; down-regulated genes and unchanged genes are indicated in blue and grey, respectively. (B) Venn diagrams showing overlap between differentially expressed genes after treatment with PAK4 siRNA or the PAK4 inhibitor. Genes that were similarly affected by PAK4 knockdown and PAK4 inhibition in VM-CUB1 cells are depicted in a heatmap generated using “ggplots” R package.

### PAK4 inhibition and PAK4 knockdown induces increased expression of PTK6

On PAK4 inhibition and knockdown, there was greater expression of PTK6, an oncogene (27). With additional treatment with a PTK6 inhibitor (dasatinib) after PAK4 inhibition, there was less cell proliferation as compared to inhibition with PAK4 or PTK6 alone **(Figure 7A**,**B)**.

**Figure 7:**
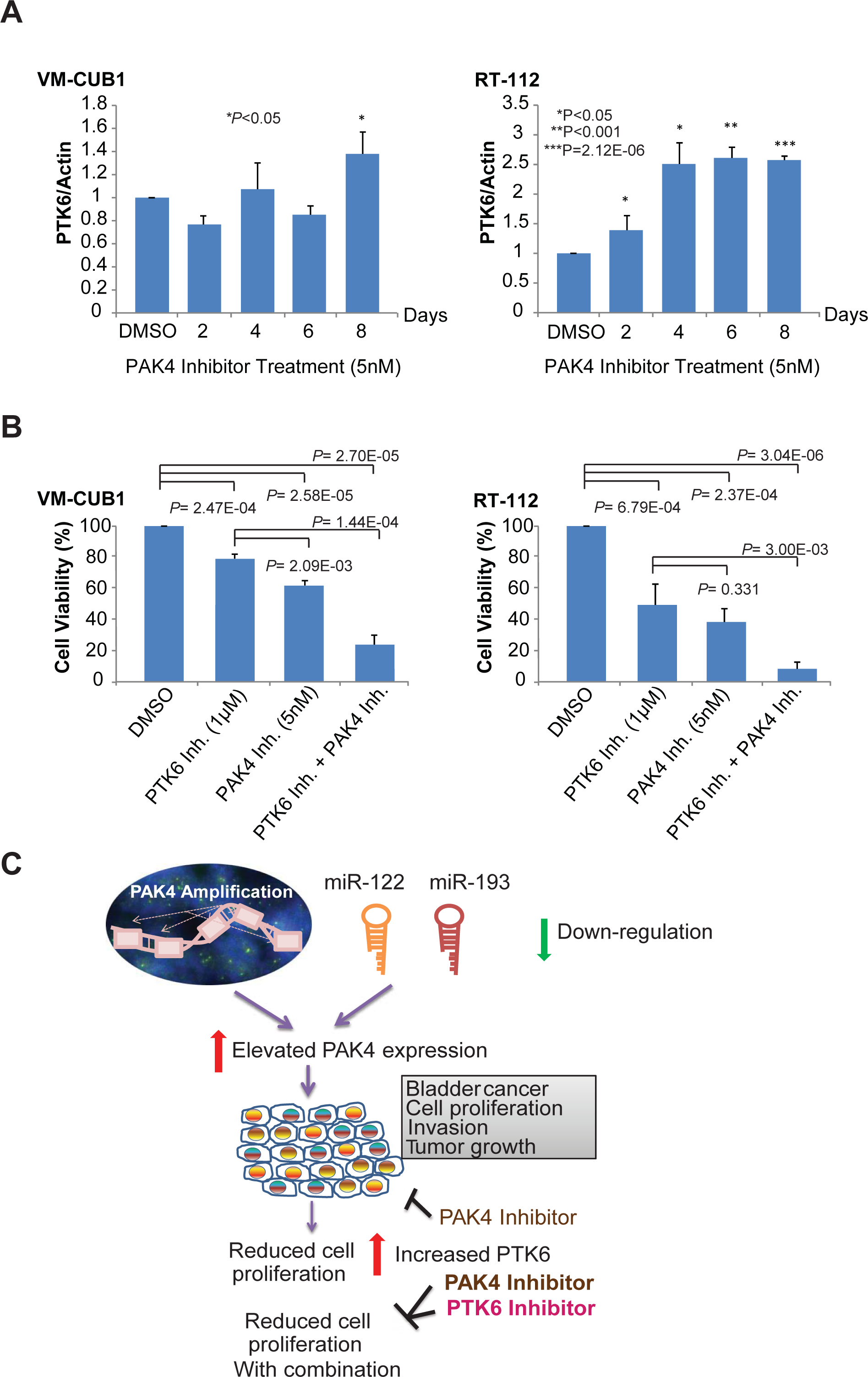
Inhibition of PTK6 following its induction induction by PAK4 inhibition results in a more extensive reduction in cell viability. (A) qRT-PCR of RNA from BLCA cells treated with 5 nM PAK4 inhibitor for a maximum of 8 days. (B) Cell viability analysis of BLCA cells treated with PTK6 inhibitor (dasatinib) and PAK4 inhibitor, alone and in combination. DMSO was used as a vehicle control. Data shown are those after 6 days of treatment. (C) Proposed model of the PAK4, miR-122/193, PTK6 regulatory axis in BLCA progression. PAK4 gene amplification and down-regulation of target miRs (miR-122, miR-193) leads to BLCA cell growth, proliferation, and invasion via elevated PAK4 expression. PAK4 inhibition leads to reduced cell proliferation, but elevates PTK6, an oncogene. Thus, combined treatment of PAK4 and PTK6 inhibitors leads to a more extensive reduction of cell proliferation.

### IPA analysis of differentially expressed genes after PAK4 inhibition and PAK4 knockdown in BLCA cells

With IPA software, canonical pathway analysis of DEGs in VM-CUB1 cells after PAK4 inhibition showed that various signaling pathways were dysregulated, including EIF2 Signaling[-log(p-value) = 9.52], Regulation of eIF4 and p70S6K Signaling [-log(p-value) = 5.08], mTOR Signaling[-log(p-value) = 4.71], Protein Kinase A Signaling[-log(p-value) = 2.85], and Integrin Signaling[-log(p-value) = 2.58] (**Supplementary Figure 4A**). Similarly, within DEGs on PAK4 knockdown, Granzyme A Signaling[-log(p-value) = 6.5], Protein Kinase A Signaling[-log(p-value) = 3.37], Cell Cycle Control of Chromosomal Replication[-log(p-value) = 3.27], Neuregulin Signaling[-log(p-value) = 3.07], and Mitotic Roles of Polo-Like Kinase[-log(p-value) = 3.06] were the top five enriched canonical pathways (**Supplementary Figure 4B**).

IPA upstream regulator analysis of DEGs after PAK4 inhibition in VM-CUB1 cells predicted an inhibitory state of EZH2 [Enhancer of Zeste homologue2] (**Supplementary Table 3**). Genes up-regulated by EZH2 (C3, CXCL8, CYP1B1, BIRC3, SLC1A3, PCDHB5, NFKBIA, MPZL2, and LCN2) were down-regulated; DKK1, which is down-regulated by EZH2, showed over-expression (**Supplementary Figure 2B**).

### Protein-protein interaction analysis reveals primary interactors affected by PAK4 inhibition/knockdown

Among the 620 differentially expressed genes in both/either of the treatments, 331 proteins were found to be interacting with 1122 connections with a confidence threshold of 0.5. The core interaction network included 277 proteins with 1085 edges (**Supplementary Figure 5**). PAK4 may directly interact (primary interactors) with six proteins (LAMA3, LAMB3, LAMC2, MAP2K1, SH3PXD2A, and RHOD). Among these, only SH3PXD2A was up-regulated upon siRNA-based knockdown of PAK4, whereas the other five were up-regulated by treatment with the inhibitor. Of these six, PAK4 interactions with three laminin proteins (LAMA3, LAMB3, and LAMC2) had a high confidence of 0.9. Five of the primary interactors (LAMA3, LAMB3, LAMC2, SH3PXD2A, and RHOD) were mainly involved in cell junction assembly. The results showed 17 proteins interacting with the six primary interactors. Most of these directly or indirectly interacting proteins were cell adhesion molecules. Except for CDK1, DEPDC1B, and ITGB8, all the primary and secondary interactors were up-regulated in response to one or both treatments. CDK1 and DEPDC1B were down-regulated by siRNA-based knockdown, whereas ITGB8 showed contradicting regulation, i.e., up-regulated by siRNA-based knockdown and down-regulated by inhibitor treatment. Manual inspection of the network showed four distinct protein hubs. Hub1 contained 19 proteins, of which 18 were significantly (q=6.80E-23) associated with translation processes. Hub2 was dominated by histone proteins representing the DNA packaging process (q=8.237E-39). The third hub of 32 proteins was enriched for cell cycle process (q=6.642E-26). Hub4 contained 20 proteins; the DNA repair process was enriched (q=7.34E-16). These *in silico* analyses provide a rationale to test some of the potential additional network genes to evaluate their role in PAK4-mediated activity in bladder cancer.

## Discussion

Identification of additional targets for therapy of MIBC is essential. In the present study, we explored the kinome of BLCA using various omic technologies including kinomic, NanoString, and RNA-sequencing. Our study identified PAK4 as amplified and overexpressed in up to 12% of bladder cancers. Using various methods, we confirmed amplification and overexpression of PAK4 with high PAK4-related kinomic activity. Furthermore, our results suggested a role for PAK4 in BLCA progression; targeting it in BLCA cells reduced cell proliferation. Thus, BLCAs often present with PAK4 overexpression, and inhibition of PAK4 reduces proliferation, colony formation, and invasion of BLCA cells.

PAK4 belongs to a family of serine/threonine kinases that are involved in cell proliferation and migration and in cytoskeletal organization. PAK4 is amplified/over-expressed in non-Hodgkin lymphoma (28), esophageal cancer (29, 30), pancreatic cancer (31-35), breast cancer (36, 37), and other cancers (38-47). Activated PAK4 a) promotes tumorigenesis in breast cancer via activation of the PI3K/AKT signaling pathway (48), b) promotes liver metastasis by phosphorylation of p53 (49), and c) facilitates the epithelial-mesenchymal transition [EMT] in prostate cancer via phosphorylation of the Slug transcription factor (50). Elevated nuclear localization of PAK4 influences breast-to-bone metastasis by targeting the metastasis suppressor, LIFR (51). Targeting of PAK4 via miRNAs, including miR-199a/b-3p (30, 39, 40, 44), miR-485 (41), miR-342 (52), miR-145 (53, 54), miR-24-1-5p (55), miR-224 (56), miR-126 (44), and miR-433 (57) suppresses cancer cell proliferation and migration.

Although BLCA, particularly MIBC, is treated with perioperative cisplatin-based combination chemotherapy, the improvement in survival is modest (3, 4). Within 2 years, systemic dissemination occurs in ∼50% of all MIBC patients. Metastatic urothelial carcinoma is generally incurable with cisplatin-based chemotherapy, yielding a median survival of 12 to 15 months. Although PD1/PD-L1 inhibitors provide durable responses for 15 to 25% of patients, the median survival of patients receiving cisplatin-based chemotherapy is less than one year (8). Given the molecular heterogeneity of the disease, rational drug development targeting molecular subgroups of patients is warranted. FGFR inhibitors have activity for patients with FGFR3-altered metastatic urothelial carcinomas, although, to yield durable benefits, an additional drug, potentially a PI3K inhibitor, may be required (58). Similarly, melanomas with BRAF-activating mutations demonstrate sensitivity to a combination of RAK and MEK inhibitors (59). The present studies showed that inhibition of PAK4, particularly in combination with a PTK6 inhibitor, is an effective therapy for the sub-set of BLCAs with PAK4 amplification or overexpression. Moreover, based on gene expression, the luminal TCGA intrinsic subtype I was enriched for PAK4 amplification and may provide a surrogate predictive marker for benefit from inhibiting PAK4.

Kinase activity profiling identified BLCAs with high PAK4 activity scores despite not having amplification. This indicates that, since kinases are frequently regulated post-transcriptionally and are activated independently, therapeutic target identification should be assessed at various biological levels (i.e., CNVs, kinase activity). PAK4 alterations occurred in 12.5-33% of the analyzed MIBCs.

We have shown that bladder cancer presents with overexpression of PAK4 and that inhibition of PAK4 inhibits the cancer phenotypes. Since many drugs engender resistance to treatments, it is necessary to assess possible downstream effects that may be induced by a therapeutic. Here, by performing sequencing of RNA from PAK4-inhibited cells, we identified activation of PTK6, a non-receptor tyrosine kinase that is overexpressed in BLCA and contributes to a poor prognosis (27). Furthermore, treatment with a PTK6 inhibitor after PAK4 inhibition caused a reduction in cell proliferation as compared to PAK4 or PTK6 inhibition alone. Together, the data support a potential therapeutic advantage of combination therapy with PAK4 and PTK6 inhibitors for patients with BLCAs overexpressing PAK4.

In summary, the present results provide evidence for overexpression of PAK4 in a subset of MIBCs. Further, PAK4 inhibition leads to upregulation of PTK6, and targeting both of these kinases reduces BLCA cell growth (**Figure 7C**). Future studies will involve use of PAK4-overexpressing BLCA patient-derived xenografts to evaluate the efficacy of PAK4 and PTK6 inhibitors in reducing BLCA growth. For selected patients, further preclinical validation and clinical trials are warranted for this potentially actionable therapeutic target.

## Material and Methods

### Patient and tumor selection

Fresh-frozen MIBC (≥pT2 stage) tissue samples with adjacent normal tissue were obtained from the Cooperative Human Tissue Network (CHTN) based at the University of Alabama at Birmingham (UAB). CHTN complies with federal human subjects regulations (The “Common Rule;” 45 CFR part 46) to collect and distribute biospecimens. Tumor samples, obtained from patients undergoing radical cystectomy without preceding neoadjuvant systemic therapy, were snap-frozen and stored in liquid N_2_ tanks (60). Specimens underwent pathological assessment for confirmation of the diagnosis. Macrodissection of tissues was conducted after histologic demarcation of tumor and normal bladder epithelial tissue. The study (X120917005) was approved by the Institutional Review Board at UAB.

### Harvesting of DNA and RNA from tissue

Tumors and normal tissues were isolated to provide genomic DNA and RNA. Genomic DNA was isolated using phenol-chloroform extraction by standard techniques and quantified by Qubit Broad Range dsDNA kits (Invitrogen, Carlsbad, CA). DNA quality was assessed by gel electrophoresis. RNA was harvested by employing Qiagen RNAeasy kits, and its quality was evaluated by the 260/280 ratio using NanoDrop (Thermo Fisher Scientific).

### Targeted kinome sequencing and data analysis

Next generation sequencing of 515 kinase genes was performed by Agilent Kinome capture and run on the Illumina MiSeq at PE150bp. Raw paired reads of length 35-151bp were aligned to the hg38 reference genome using the BWA-MEM algorithm of the Burrows-Wheeler Aligner (V 0.7.12) with default settings (61). The BWA-aligned reads were further processed as per recommendations from GATK best practices. Picard suite (V 1.140) [http://broadinstitute.github.io/picard/index.html] was used to 1) sort and convert sam files to bam [SortSam], 2) mark duplicate reads to ignore from next analysis steps [MarkDuplicate], and 3) generate bam index files [BuildBamIndex]. Further, each bam file was subjected to indel realignment and base quality recalibration using Genome Analysis Toolkit (GATK) V 3.5 (62, 63). Subsequently, somatic single nucleotide polymorphisms (SNPs) and somatic insertion/deletions (INDELs) were screened within kinases of each BLCA sample in comparison with their matched normal tissue. Somatic mutations were identified independently with MuTect2 (64) and Varscan2 (65).

The somatic mutations were filtered and annotated using ANNOVAR (66). The somatic mutations that were a) found in non-exonic regions; b) already reported in dbSNP 144 (67) and not reported in COSMIC 70 (68) databases; and c) leading to synonymous amino acid changes, were filtered out. Thus, 183 and 168 somatic SNVs/INDELs were identified by MuTect and Varscan2 respectively. In total, 127 somatic SNVs/INDELs in the human kinome were commonly identified by both tools.

Varscan2 was used to identify somatic copy number alterations in each BLCA by comparing with matched normal tissue. Using mpileup from Samtools (69) and the copy number method from varscan2, raw copy number calls were obtained from BAM files for each normal and tumor pair. Next, the Copycaller method from Varscan2 was used to adjust raw copy number calls to GC content. This was followed by application of circular binary segmentation using R package ‘DNAcopy’. Finally, adjacent segments with similar copy numbers were merged and classified based on size (large-scale or focal). Circos plots depicting results of kinome sequencing analysis were generated using Circos software (70). Targeted kinome sequencing data have been submitted to the NCBI Sequence Read Archive (SRA) [Accession ID: PRJNA548509].

### Kinase gene expression assay on the NanoString platform

Expression profiling of 519 kinase genes and 8 housekeeping genes was measured using the NanoString nCounter® analysis system (71). RNA was hybridized for 19 hours at 65°C in the nCounter® NanoString platform, followed by digital counting utilizing two hybridizing base probes per RNA. A codeset specific to a 100-base region per RNA, which used a 3’ biotinylated capture probe and a 5’ reporter probe tagged with a fluorescent barcode, was employed. Background hybridization was assessed by spiked-in negative controls. Normalization and differential expression analyses were performed using the advanced analysis plugin of nSolver software (NanoString Technologies). Differential expression was assessed considering normalized digital raw counts of 24 tumor and 20 matched normal samples. Four matched normal samples flagged during normalization were not considered for differential expression analysis. Genes with absolute fold changes of >=1.5 or <= −1.5 and P values <0.05 were considered as differentially expressed. The Nanostring data have been deposited at NCBI Gene Expression Omnibus [GEO, #GSE130598].

### Kinase activity profiling utilizing PamStation microarray

Lysates of paired normal tissues and tumors from 24 BLCA patients were analyzed for enzymatic kinase activity on PamChips (PamGene, Den Bosch, Netherlands) in the UAB Kinome Core (www.kinomecore.com) (72, 73). Briefly, lysates were repeatedly pumped through PamChip 3D microarrays containing ∼288 phosphorylatable 12-15 amino acid tyrosine-, serine-, or threonine-kinase substrates, and phosphorylation intensity as measured by phospho-specific FITC-conjugated antibodies was captured over time in a computer controlled manner. Both kinetic and end-level phosphorylation levels were captured. Log2 transformed end-level values were used for comparison. We compared the phosphorylation of four PAK4 peptides, CFTR_730_742, NMDZ1_890_902, KS6A1_374_386, and TOP2A_1463_1475, which were identified via Kinexus database as ranking PAK4 in the top 20 listed kinases per each peptide (**Figure 3A**). For each peptide, the ranking of PAK4 at the time of analysis was as follows: CFTR_730_742 (11^th^), NMDZ1_890_902 (12^th^), KS6A1_374_386 (14^th^), and TOP2A_1463_1475 (14^th^). PAK4 kinase activity scoring (PKAS) was measured as subtracted Log2 signal (tumor minus normal) and dividing by per-peptide rank, and taking the mean across four peptides. PKAS values > 2.0 were considered high.

### *In silico* validation using the BLCA TCGA dataset

Using cBioPortal.org (74), mutation and copy number status profiles of most frequently mutated/altered kinases were examined in bladder urothelial carcinoma (BLCA) data from provisional TCGA. The analysis results were downloaded as OncoPrint. Survival data for TCGA BLCA patients (n=412) with/without PAK4 alterations [amplification or mRNA dysregulation] was obtained from cBioPortal. UALCAN (75) was used to obtain PAK4 expression profiles based on TCGA level 3 RNA-seq data for BLCAs (n=408) and adjacent normal samples (n=19).

### RNA-seq data analysis

Raw sequencing data comprising 50bp paired-end reads were cleaned using Trim Galore (v0.4.1) [http://www.bioinformatics.babraham.ac.uk/projects/trim_galore/] and subjected to quality control using FastQC (v0.11.5) [http://www.bioinformatics.babraham.ac.uk/projects/fastqc/]. The quality trimmed reads were aligned to human genome (GRCh38) using TopHat v2.1.0 (76). The aligned reads were sorted using Samtools (Version: 1.3.1) (77), and, by use of HTSeq-count (version 0.6.0) (78), the numbers of reads aligned to each annotated human gene were enumerated. Finally, an R package “DESeq” (79) was used to normalize raw read counts and to perform differentialexpressionanalysisfollowingastandardprotocol [https://bioconductor.org/packages/release/bioc/vignettes/DESeq/inst/doc/DESeq.pdf]. Genes with absolute fold changes of ≥1.5 and p-values <0.05 were selected as differentially expressed genes (DEGs). Venny [http://bioinfogp.cnb.csic.es/tools/venny/index.html] was used to obtain common genes differentially expressed in BLCA VM-CUB1 cells after treatment with PAK4 siRNA or the PAK4 inhibitor. Heatmaps of the top differentially expressed genes were created using the heatmap.2 function of R package ‘gplots’ [http://CRAN.R-project.org/package=gplots]. RNA-seq data were uploaded to the NCBI Gene Expression Omnibus [GEO, #GSE130455].

### Pathway and upstream regulator analysis

In VM-CUB1 cells after PAK4 inhibitor or PAK4 siRNA treatment, canonical pathways enriched by DEGs were identified separately using the Ingenuity Pathway Analysis (IPA) Core analysis module. Similarly, IPA upstream regulator analysis was used to identify potential upstream regulators of DEGs.

### Cell culture

The bladder cancer cell lines HT1197 (ATCC), HT1376 (ATCC), VM-CUB1 (DSMZ), and KU-19-19 (DSMZ) were grown in 90% MEM (Corning™, NY) + 10% fetal bovine serum (FBS, Invitrogen, Thermo Fisher Scientific, Carlsbad, CA). The BLCA cell lines T24 (ATCC), 5637 (ATCC), and RT-112 (DSMZ) were grown in 90% RPMI 1640 (Life Technologies, CA) + 10% FBS with 100 U/ml penicillin-streptomycin in a 5% CO_2_ cell culture incubator.

### Benign and Tumor Tissues

As described earlier (80), we utilized formalin-fixed paraffin-embedded tissues, both normal tissues and clinically localized BLCAs. **Supplementary Table 4** provides demographics and clinical characteristics of patients considered in the study. The bladder tissues were collected in a retrospective study approved by the Institutional Review Board at the University of Alabama at Birmingham (UAB), which allowed the investigation of de-identified samples obtained from human subjects.

### siRNA and miRNA Transfections

Small interfering RNA (siRNA) duplexes targeting PAK4, PAK4 siRNA 1 (D-003615-06-0020) and 2 (D-003615-07-0020), and non-targeting (NT) siRNA were purchased from Dharmacon (Lafayette, CO). Precursors of respective human microRNAs (miR-27a, −122, −128, −193 and −217) and negative controls were purchased from Life Technologies, Thermo Fisher Scientific, CA. Transfections were performed with Lipofectamine RNAiMAX Reagent (Life Technologies). For si/miRNA transfections, VM-CUB1 and RT-112 cells were seeded at 1 × 10^5^ cells per well in 6-well plates along with siRNA duplexes or miRNAs using Lipofectamine RNAiMAX Reagent (Life Technologies). At 72 hours after transfection, cells were either seeded for cell proliferation, colony formation, and invasion assays or were harvested for RNA isolation or immunoblot analysis.

### Cell Proliferation

Cell proliferation assays were conducted for siRNA-treated cells and measured by cell counting. Cells with transient or stable PAK4 knock down were plated at 10,000 cells/well in 12-well plates (*n* = 3). Cells were harvested and, at indicated time points, counted with a Coulter counter (Beckman Coulter, Fullerton, CA). Cells treated with NT siRNA served as controls.

### Cell Viability Assays

For inhibitor-treated cells, viability assays were conducted by using CellTiter-Glo (Promega, Madison, WI). VM-CUB1 and RT-112 cells were seeded in 96-well plates on day 0, treated with various concentrations of PAK4 inhibitor on day 1, and incubated for 2, 4, or 6 days. For combined PAK4 and PTK6 treatements, the cells were incubated for 8 days. Untreated and DMSO-treated cells served as controls. Cells were then trypsinized and seeded at a density of 250 cells per well in 96-well plates (*n*=3), then treated with a Protein Tyrosine Kinase 6 (PTK6) inhibitor (dasatinib, catalogue # NC0713371, Thermo Fisher Scientific) or PAK4 inhibitor (vandetanib, catalogue # NC0706691, Thermo Fisher Scientific), alone or in combination. CellTiter-Glo was added, and luminescence was measured at specified time points. Experiments were performed with three replicates per sample.

### Real-Time Quantitative PCR

RNA was isolated from normal bladder tissues and from VM-CUB1 and RT-112 cells using Direct-zol RNA MiniPrep kits (Zymo Research, Irvine, CA). Next, RNA was reverse transcribed into complementary DNA using Superscript III Reverse Transcriptase (Invitrogen). SYBR green real-time quantitative PCR (qRT-PCR) analysis was performed using primers for ACTB, PAK4, or PTK6. The oligonucleotide sequences were as follows: ACTB-F (5’-3’) GCACAGAGCCTCGCCTT; ACTB-R (5’-3’) GTTGTCGACGACGAGCG; PAK4-F (5’-3’) GTGCAAGAGAGCTGAGGGAG; PAK4-R (5’-3’) CTCTAGGGGCTTCGGGTTAC; PTK6-F (5’-3’) GGCTATGTGCCCCACAACTA; and PTK6-R (5’-3’) GACGCACAGCTTCCGAG. Samples were tested in triplicate.

### Western Blot Analyses

Immunoblot analyses were performed as described earlier (80). Briefly, cell lysates were prepared in NP-40 lysis buffer (Boston Bioproducts, Ashland, MA) with 1X Halt protease inhibitor cocktail (Thermo Fisher Scientific, Grand Island, NY). Protein was quantified by use of BioRad DC protein assays (Bio-Rad Laboratories, Hercules, CA), and 10-μg protein samples were separated on NuPAGE 4%-12% Bis-Tris protein gels and transferred onto Immobilon-P PVDF membranes (EMD Millipore, Billerica, MA). The membranes were incubated for 1 hour in blocking buffer (Tris-buffered saline with 0.1% Tween and 5% nonfat dry milk) followed by overnight incubation at 4°C with the primary antibody. Then the blots were incubated with horseradish peroxidase– conjugated secondary antibody (1:5000) for 1 hour at room temperature, and signals were visualized by Luminata Forte chemiluminescence Western blotting substrate as per the manufacturer’s protocol (EMD Millipore). Antibodies used were rabbit anti-PAK4 #3242 (IB, 1:1000; Cell Signaling Technology, Danvers, MA) and anti–HRP-β-actin: # HRP-60008 (IB, 1:200000; PTG Labs, Rosemont, IL).

### Matrigel Invasion Assays

Matrigel invasion assays were performed as described earlier (80-82). Briefly, cells were seeded onto Corning BioCoat Matrigel matrices (# 08-774-122, Corning, New York, NY) in the upper chambers of 24-well culture plates without FBS. The lower chambers contained the respective medium supplemented with 10% FBS as a chemoattractant. After 48 hours, the non-invading cells and the Matrigel matrices were removed with cotton swabs. Invasive cells located on the lower sides of the chambers were stained with 0.2% crystal violet in methanol, air-dried, and photographed using an inverted microscope (4×). Invasion was quantified by a colorimetric assay. For this, the inserts were treated with 150 μl of 10% acetic acid, and absorbance was measured at 560 nm.

### Colony Formation Assays

Colony formation assays were performed as described earlier (80). Transient, stable knockdown, or inhibitor-treated cells were counted and seeded as 1000 cells per well of 6-well plates (triplicates) and incubated at 37°C, with 5% CO_2_, for 10-15 days. Colonies were fixed with 10% (v/v) glutaraldehyde for 30 minutes and stained with crystal violet (Sigma-Aldrich, St. Louis, MO) for 20 minutes. Then, photographs of the colonies were taken using an Amersham Imager 600RGB (GE Healthcare Life Sciences, Pittsburgh, PA).

### Protein-protein interaction network analysis

The differentially expressed genes obtained by sequencing of RNA for VM-CUB1 cells treated with PAK4 inhibitor or siRNA against PAK4 were used for generation of protein interaction networks. The protein interactions among differential genes were retrieved from the STRING database (83). The search was restricted to interactions with ‘experimental’ evidence and from ‘databases’, and a minimum confidence threshold of 0.5 was applied. The obtained interactions were then submitted to Cytoscape network analysis (84) for visualization and analysis. The protein interaction network was manually inspected to extract the hubs of densely interacting proteins. The functional analysis and enrichment of the protein-protein interaction network was performed using the KEGG database (85), ToppGene Suite (86), and TRRUST database (87).

### Statistical analyses

Coefficients of drug interaction (CDIs) were calculated with the equation CDI = AB/(A × B). Briefly, the relative cell viability of the combination (AB) was divided by the relative cell viabilities of the single agents multiplied. CDI < 1 indicated a synergistic effect; CDI = 1 indicated an additive effect; and CDI > 1 indicated an antagonistic effect. This calculation was performed for each set of drug concentrations (88).

## Funding /acknowledgement

This study was supported in part by institutional funds (Department of Pathology and School of Medicine of the University of Alabama at Birmingham) awarded to to S.V. IBAB is supported by the Department of IT, BT and S&T, Government of Karnataka, India. The authors thank Dr. Donald Hill for critical reading and editing of this manuscript.

## References

1. Antoni S, Ferlay J, Soerjomataram I, Znaor A, Jemal A, Bray F. Bladder Cancer Incidence and Mortality: A Global Overview and Recent Trends. Eur Urol. 2017;71(1):96–108.

2. von der Maase H, Sengelov L, Roberts JT, Ricci S, Dogliotti L, Oliver T, et al. Long-term survival results of a randomized trial comparing gemcitabine plus cisplatin, with methotrexate, vinblastine, doxorubicin, plus cisplatin in patients with bladder cancer. J Clin Oncol. 2005;23(21):4602–8.

3. Grossman HB, Natale RB, Tangen CM, Speights VO, Vogelzang NJ, Trump DL, et al. Neoadjuvant chemotherapy plus cystectomy compared with cystectomy alone for locally advanced bladder cancer. The New England journal of medicine. 2003;349(9):859–66.

4. Griffiths G, Hall R, Sylvester R, Raghavan D, Parmar MK. International phase III trial assessing neoadjuvant cisplatin, methotrexate, and vinblastine chemotherapy for muscle-invasive bladder cancer: long-term results of the BA06 30894 trial. Journal of clinical oncology : official journal of the American Society of Clinical Oncology. 2011;29(16):2171–7.

5. Rosenberg JE, Hoffman-Censits J, Powles T, van der Heijden MS, Balar AV, Necchi A, et al. Atezolizumab in patients with locally advanced and metastatic urothelial carcinoma who have progressed following treatment with platinum-based chemotherapy: a single-arm, multicentre, phase 2 trial. Lancet. 2016;387(10031):1909–20.

6. Bellmunt J, de Wit R, Vaughn DJ, Fradet Y, Lee JL, Fong L, et al. Pembrolizumab as Second-Line Therapy for Advanced Urothelial Carcinoma. N Engl J Med. 2017;376(11):1015–26.

7. Sharma P, Callahan MK, Bono P, Kim J, Spiliopoulou P, Calvo E, et al. Nivolumab monotherapy in recurrent metastatic urothelial carcinoma (CheckMate 032): a multicentre, open-label, two-stage, multiarm, phase 1/2 trial. Lancet Oncol. 2016;17(11):1590–8.

8. Sonpavde G. PD-1 and PD-L1 Inhibitors as Salvage Therapy for Urothelial Carcinoma. N Engl J Med. 2017;376(11):1073–4.

9. Balar AV, Galsky MD, Rosenberg JE, Powles T, Petrylak DP, Bellmunt J, et al. Atezolizumab as first-line treatment in cisplatin-ineligible patients with locally advanced and metastatic urothelial carcinoma: a single-arm, multicentre, phase 2 trial. Lancet. 2017;389(10064):67–76.

10. Balar AV, Castellano D, O’Donnell PH, Grivas P, Vuky J, Powles T, et al. First-line pembrolizumab in cisplatin-ineligible patients with locally advanced and unresectable or metastatic urothelial cancer (KEYNOTE-052): a multicentre, single-arm, phase 2 study. Lancet Oncol. 2017.

11. Choi W, Porten S, Kim S, Willis D, Plimack ER, Hoffman-Censits J, et al. Identification of distinct Basal and luminal subtypes of muscle-invasive bladder cancer with different sensitivities to frontline chemotherapy. Cancer Cell. 2014;25(2):152–65.

12. McConkey DJ, Choi W, Ochoa A, Siefker-Radtke A, Czerniak B, Dinney CP. Therapeutic opportunities in the intrinsic subtypes of muscle-invasive bladder cancer. Hematol Oncol Clin North Am. 2015;29(2):377-94, x-xi.

13. Iyer G, Al-Ahmadie H, Schultz N, Hanrahan AJ, Ostrovnaya I, Balar AV, et al. Prevalence and cooccurrence of actionable genomic alterations in high-grade bladder cancer. J Clin Oncol. 2013;31(25):3133–40.

14. Robertson AG, Kim J, Al-Ahmadie H, Bellmunt J, Guo G, Cherniack AD, et al. Comprehensive Molecular Characterization of Muscle-Invasive Bladder Cancer. Cell. 2017;171(3):540–56 e25.

15. Jacobs JJ, van Lohuizen M. Cellular memory of transcriptional states by Polycomb-group proteins. Semin Cell Dev Biol. 1999;10(2):227–35.

16. Francis NJ, Kingston RE. Mechanisms of transcriptional memory. Nat Rev Mol Cell Biol. 2001;2(6):409–21.

17. Sonpavde G, Jones BS, Bellmunt J, Choueiri TK, Sternberg CN. Future directions and targeted therapies in bladder cancer. Hematol Oncol Clin North Am. 2015;29(2):361-76, x.

18. Bellmunt J, Orsola A, Sonpavde G. Precision and Predictive Medicine in Urothelial Cancer: Are We Making Progress? Eur Urol. 2015.

19. Erdafitinib Efficacious in Bladder Cancer. Cancer Discov. 2018;8(8):OF6.

20. Choudhury NJ, Campanile A, Antic T, Yap KL, Fitzpatrick CA, Wade JL, 3rd, et al. Afatinib Activity in Platinum-Refractory Metastatic Urothelial Carcinoma in Patients With ERBB Alterations. J Clin Oncol. 2016;34(18):2165–71.

21. Iyer G, Hanrahan AJ, Milowsky MI, Al-Ahmadie H, Scott SN, Janakiraman M, et al. Genome sequencing identifies a basis for everolimus sensitivity. Science. 2012;338(6104):221.

22. Agarwal V, Bell GW, Nam JW, Bartel DP. Predicting effective microRNA target sites in mammalian mRNAs. Elife. 2015;4.

23. Bandiera S, Pfeffer S, Baumert TF, Zeisel MB. miR-122--a key factor and therapeutic target in liver disease. J Hepatol. 2015;62(2):448–57.

24. Jian B, Li Z, Xiao D, He G, Bai L, Yang Q. Downregulation of microRNA-193-3p inhibits tumor proliferation migration and chemoresistance in human gastric cancer by regulating PTEN gene. Tumour Biol. 2016;37(7):8941–9.

25. Zhang HS, Zhang FJ, Li H, Liu Y, Du GY, Huang YH. Tanshinone A inhibits human esophageal cancer cell growth through miR-122-mediated PKM2 down-regulation. Arch Biochem Biophys. 2016;598:50–6.

26. Wang Y, Xing QF, Liu XQ, Guo ZJ, Li CY, Sun G. MiR-122 targets VEGFC in bladder cancer to inhibit tumor growth and angiogenesis. Am J Transl Res. 2016;8(7):3056–66.

27. Xu XL, Ye YL, Wu ZM, He QM, Tan L, Xiao KH, et al. Overexpression of PTK6 predicts poor prognosis in bladder cancer patients. J Cancer. 2017;8(17):3464–73.

28. Li N, Lopez MA, Linares M, Kumar S, Oliva S, Martinez-Lopez J, et al. Dual PAK4-NAMPT inhibition impacts growth and survival, and increases sensitivity to DNA-damaging agents in Waldenstrom Macroglobulinemia. Clin Cancer Res. 2018.

29. Jiang YY, Lin DC, Mayakonda A, Hazawa M, Ding LW, Chien WW, et al. Targeting super-enhancer-associated oncogenes in oesophageal squamous cell carcinoma. Gut. 2017;66(8):1358–68.

30. Phatak P, Burrows WM, Chesnick IE, Tulapurkar ME, Rao JN, Turner DJ, et al. MiR-199a-3p decreases esophageal cancer cell proliferation by targeting p21 activated kinase 4. Oncotarget. 2018;9(47):28391–407.

31. Mahlamaki EH, Kauraniemi P, Monni O, Wolf M, Hautaniemi S, Kallioniemi A. High-resolution genomic and expression profiling reveals 105 putative amplification target genes in pancreatic cancer. Neoplasia. 2004;6(5):432–9.

32. Chen S, Auletta T, Dovirak O, Hutter C, Kuntz K, El-ftesi S, et al. Copy number alterations in pancreatic cancer identify recurrent PAK4 amplification. Cancer Biol Ther. 2008;7(11):1793–802.

33. Kimmelman AC, Hezel AF, Aguirre AJ, Zheng H, Paik JH, Ying H, et al. Genomic alterations link Rho family of GTPases to the highly invasive phenotype of pancreas cancer. Proc Natl Acad Sci U S A. 2008;105(49):19372–7.

34. Tyagi N, Bhardwaj A, Singh AP, McClellan S, Carter JE, Singh S. p-21 activated kinase 4 promotes proliferation and survival of pancreatic cancer cells through AKT- and ERK-dependent activation of NF-kappaB pathway. Oncotarget. 2014;5(18):8778–89.

35. Thillai K, Sarker D, Wells C. PAK4 pathway as a potential therapeutic target in pancreatic cancer. Future Oncol. 2018;14(7):579–82.

36. Yu W, Kanaan Y, Bae YK, Gabrielson E. Chromosomal changes in aggressive breast cancers with basal-like features. Cancer Genet Cytogenet. 2009;193(1):29–37.

37. Minden A. The pak4 protein kinase in breast cancer. ISRN Oncol. 2012;2012:694201.

38. Park MH, Lee HS, Lee CS, You ST, Kim DJ, Park BH, et al. p21-Activated kinase 4 promotes prostate cancer progression through CREB. Oncogene. 2013;32(19):2475–82.

39. Callegari E, D’Abundo L, Guerriero P, Simioni C, Elamin BK, Russo M, et al. miR-199a-3p Modulates MTOR and PAK4 Pathways and Inhibits Tumor Growth in a Hepatocellular Carcinoma Transgenic Mouse Model. Mol Ther Nucleic Acids. 2018;11:485–93.

40. Zeng B, Shi W, Tan G. MiR-199a/b-3p inhibits gastric cancer cell proliferation via downregulating PAK4/MEK/ERK signaling pathway. BMC Cancer. 2018;18(1):34.

41. Mao K, Lei D, Zhang H, You C. MicroRNA-485 inhibits malignant biological behaviour of glioblastoma cells by directly targeting PAK4. Int J Oncol. 2017;51(5):1521–32.

42. Begum A, Imoto I, Kozaki K, Tsuda H, Suzuki E, Amagasa T, et al. Identification of PAK4 as a putative target gene for amplification within 19q13.12-q13.2 in oral squamous-cell carcinoma. Cancer Sci. 2009;100(10):1908–16.

43. Davis SJ, Sheppard KE, Pearson RB, Campbell IG, Gorringe KL, Simpson KJ. Functional analysis of genes in regions commonly amplified in high-grade serous and endometrioid ovarian cancer. Clin Cancer Res. 2013;19(6):1411–21.

44. Li SQ, Wang ZH, Mi XG, Liu L, Tan Y. MiR-199a/b-3p suppresses migration and invasion of breast cancer cells by downregulating PAK4/MEK/ERK signaling pathway. IUBMB Life. 2015;67(10):768–77.

45. Cai S, Ye Z, Wang X, Pan Y, Weng Y, Lao S, et al. Overexpression of P21-activated kinase 4 is associated with poor prognosis in non-small cell lung cancer and promotes migration and invasion. J Exp Clin Cancer Res. 2015;34:48.

46. Liu W, Yang Y, Liu Y, Liu H, Zhang W, Xu L, et al. p21-Activated kinase 4 predicts early recurrence and poor survival in patients with nonmetastatic clear cell renal cell carcinoma. Urol Oncol. 2015;33(5):205 e13–21.

47. Song B, Wang W, Zheng Y, Yang J, Xu Z. P21-activated kinase 1 and 4 were associated with colorectal cancer metastasis and infiltration. J Surg Res. 2015;196(1):130–5.

48. He LF, Xu HW, Chen M, Xian ZR, Wen XF, Chen MN, et al. Activated-PAK4 predicts worse prognosis in breast cancer and promotes tumorigenesis through activation of PI3K/AKT signaling. Oncotarget. 2017;8(11):17573–85.

49. Xu HT, Lai WL, Liu HF, Wong LL, Ng IO, Ching YP. PAK4 Phosphorylates p53 at Serine 215 to Promote Liver Cancer Metastasis. Cancer Res. 2016;76(19):5732–42.

50. Park JJ, Park MH, Oh EH, Soung NK, Lee SJ, Jung JK, et al. The p21-activated kinase 4-Slug transcription factor axis promotes epithelial-mesenchymal transition and worsens prognosis in prostate cancer. Oncogene. 2018.

51. Li Y, Zhang H, Zhao Y, Wang C, Cheng Z, Tang L, et al. A mandatory role of nuclear PAK4-LIFR axis in breast-to-bone metastasis of ERalpha-positive breast cancer cells. Oncogene. 2018.

52. Lu X, Wang H, Su Z, Cai L, Li W. MicroRNA-342 inhibits the progression of glioma by directly targeting PAK4. Oncol Rep. 2017;38(2):1240–50.

53. Sheng N, Tan G, You W, Chen H, Gong J, Chen D, et al. MiR-145 inhibits human colorectal cancer cell migration and invasion via PAK4-dependent pathway. Cancer Med. 2017;6(6):1331–40.

54. Azmi AS, Li Y, Muqbil I, Aboukameel A, Senapedis W, Baloglu E, et al. Exportin 1 (XPO1) inhibition leads to restoration of tumor suppressor miR-145 and consequent suppression of pancreatic cancer cell proliferation and migration. Oncotarget. 2017;8(47):82144–55.

55. Zhang W, Fei J, Yu S, Shen J, Zhu X, Sadhukhan A, et al. LINC01088 inhibits tumorigenesis of ovarian epithelial cells by targeting miR-24-1-5p. Sci Rep. 2018;8(1):2876.

56. Xia M, Wei J, Tong K. MiR-224 promotes proliferation and migration of gastric cancer cells through targeting PAK4. Pharmazie. 2016;71(8):460–4.

57. Xue J, Chen LZ, Li ZZ, Hu YY, Yan SP, Liu LY. MicroRNA-433 inhibits cell proliferation in hepatocellular carcinoma by targeting p21 activated kinase (PAK4). Mol Cell Biochem. 2015;399(1-2):77–86.

58. Siefker-Radtke AO, Necchi A, Park SH, GarcÃa-Donas Js, Huddart RA, Burgess EF, et al. First results from the primary analysis population of the phase 2 study of erdafitinib (ERDA; JNJ-42756493) in patients (pts) with metastatic or unresectable urothelial carcinoma (mUC) and FGFR alterations (FGFRalt). Journal of Clinical Oncology. 2018;36(15_suppl):4503-.

59. Robert C, Karaszewska B, Schachter J, Rutkowski P, Mackiewicz A, Stroiakovski D, et al. Improved overall survival in melanoma with combined dabrafenib and trametinib. N Engl J Med. 2015;372(1):30–9.

60. Grizzle WE BW, Fredenburgh J. Safety in biomedical and other laboratories. In: Molecular Diagnostics. (Eds G Patrinos W Ansorg) Chapter 33, pp 421–428, 2005.

61. Li H, Durbin R. Fast and accurate short read alignment with Burrows-Wheeler transform. Bioinformatics. 2009;25(14):1754–60.

62. McKenna A, Hanna M, Banks E, Sivachenko A, Cibulskis K, Kernytsky A, et al. The Genome Analysis Toolkit: a MapReduce framework for analyzing next-generation DNA sequencing data. Genome Res. 2010;20(9):1297–303.

63. DePristo MA, Banks E, Poplin R, Garimella KV, Maguire JR, Hartl C, et al. A framework for variation discovery and genotyping using next-generation DNA sequencing data. Nat Genet. 2011;43(5):491–8.

64. Cibulskis K, Lawrence MS, Carter SL, Sivachenko A, Jaffe D, Sougnez C, et al. Sensitive detection of somatic point mutations in impure and heterogeneous cancer samples. Nat Biotechnol. 2013;31(3):213–9.

65. Koboldt DC, Zhang Q, Larson DE, Shen D, McLellan MD, Lin L, et al. VarScan 2: somatic mutation and copy number alteration discovery in cancer by exome sequencing. Genome Res. 2012;22(3):568–76.

66. Wang K, Li M, Hakonarson H. ANNOVAR: functional annotation of genetic variants from highthroughput sequencing data. Nucleic Acids Res. 2010;38(16):e164.

67. Sherry ST, Ward MH, Kholodov M, Baker J, Phan L, Smigielski EM, et al. dbSNP: the NCBI database of genetic variation. Nucleic Acids Res. 2001;29(1):308–11.

68. Forbes SA, Tang G, Bindal N, Bamford S, Dawson E, Cole C, et al. COSMIC (the Catalogue of Somatic Mutations in Cancer): a resource to investigate acquired mutations in human cancer. Nucleic Acids Res. 2010;38(Database issue):D652–7.

69. Li H, Handsaker B, Wysoker A, Fennell T, Ruan J, Homer N, et al. The Sequence Alignment/Map format and SAMtools. Bioinformatics. 2009;25(16):2078–9.

70. Krzywinski M, Schein J, Birol I, Connors J, Gascoyne R, Horsman D, et al. Circos: an information aesthetic for comparative genomics. Genome Res. 2009;19(9):1639–45.

71. Geiss GK, Bumgarner RE, Birditt B, Dahl T, Dowidar N, Dunaway DL, et al. Direct multiplexed measurement of gene expression with color-coded probe pairs. Nat Biotechnol. 2008;26(3):317–25.

72. Anderson JC, Willey CD, Mehta A, Welaya K, Chen D, Duarte CW, et al. High Throughput Kinomic Profiling of Human Clear Cell Renal Cell Carcinoma Identifies Kinase Activity Dependent Molecular Subtypes. PLoS One. 2015;10(9):e0139267.

73. Ghosh AP, Willey CD, Anderson JC, Welaya K, Chen D, Mehta A, et al. Kinomic profiling identifies focal adhesion kinase 1 as a therapeutic target in advanced clear cell renal cell carcinoma. Oncotarget. 2017;8(17):29220–32.

74. Gao J, Aksoy BA, Dogrusoz U, Dresdner G, Gross B, Sumer SO, et al. Integrative analysis of complex cancer genomics and clinical profiles using the cBioPortal. Sci Signal. 2013;6(269):pl1.

75. Chandrashekar DS, Bashel B, Balasubramanya SAH, Creighton CJ, Ponce-Rodriguez I, Chakravarthi B, et al. UALCAN: A Portal for Facilitating Tumor Subgroup Gene Expression and Survival Analyses. Neoplasia. 2017;19(8):649–58.

76. Kim D, Pertea G, Trapnell C, Pimentel H, Kelley R, Salzberg SL. TopHat2: accurate alignment of transcriptomes in the presence of insertions, deletions and gene fusions. Genome Biol. 2013;14(4):R36.

77. Li H. A statistical framework for SNP calling, mutation discovery, association mapping and population genetical parameter estimation from sequencing data. Bioinformatics. 2011;27(21):2987–93.

78. Anders S, Pyl PT, Huber W. HTSeq--a Python framework to work with high-throughput sequencing data. Bioinformatics. 2015;31(2):166–9.

79. Anders S, Huber W. Differential expression analysis for sequence count data. Genome Biol. 2010;11(10):R106.

80. Chakravarthi B, Rodriguez Pena MDC, Agarwal S, Chandrashekar DS, Hodigere Balasubramanya SA, Jabboure FJ, et al. A Role for De Novo Purine Metabolic Enzyme PAICS in Bladder Cancer Progression. Neoplasia. 2018;20(9):894–904.

81. Chakravarthi BV, Pathi SS, Goswami MT, Cieslik M, Zheng H, Nallasivam S, et al. The miR-124-prolyl hydroxylase P4HA1-MMP1 axis plays a critical role in prostate cancer progression. Oncotarget. 2014;5(16):6654–69.

82. Chakravarthi BV, Goswami MT, Pathi SS, Dodson M, Chandrashekar DS, Agarwal S, et al. Expression and Role of PAICS, a De Novo Purine Biosynthetic Gene in Prostate Cancer. Prostate. 2017;77(1):10–21.

83. Szklarczyk D, Morris JH, Cook H, Kuhn M, Wyder S, Simonovic M, et al. The STRING database in 2017: quality-controlled protein-protein association networks, made broadly accessible. Nucleic Acids Res. 2017;45(D1):D362–D8.

84. Shannon P, Markiel A, Ozier O, Baliga NS, Wang JT, Ramage D, et al. Cytoscape: a software environment for integrated models of biomolecular interaction networks. Genome Res. 2003;13(11):2498–504.

85. Kanehisa M, Goto S. KEGG: kyoto encyclopedia of genes and genomes. Nucleic Acids Res. 2000;28(1):27–30.

86. Chen J, Bardes EE, Aronow BJ, Jegga AG. ToppGene Suite for gene list enrichment analysis and candidate gene prioritization. Nucleic Acids Res. 2009;37(Web Server issue):W305–11.

87. Han H, Cho JW, Lee S, Yun A, Kim H, Bae D, et al. TRRUST v2: an expanded reference database of human and mouse transcriptional regulatory interactions. Nucleic Acids Res. 2018;46(D1):D380–D6.

88. Hao JQ, Li Q, Xu SP, Shen YX, Sun GY. Effect of lumiracoxib on proliferation and apoptosis of human nonsmall cell lung cancer cells in vitro. Chin Med J (Engl). 2008;121(7):602–7.

