## Supplementary Figures for "Therapeutically actionable PAK4 is amplified, overexpressed and involved in bladder cancer progression"

### Supplementary Figure legends

**Supplementary Figure 1:** (A, B) Validation of most frequently amplified and deleted kinases using TCGA Bladder urothelial carcinoma (BLCA) exome sequencing data. Oncoprint from cBioPortal shows proportion of 408 BLCA samples with copy number alterations in kinases of interest. (C-E) Boxplots showing RNA-level expression of PAK4 in TCGA BLCA and normal samples based on pathological stages, molecular subtypes and tumor histology.

**Supplementary Figure 2:** (A) Kaplan Meir plot showing impact of PAK4 amplification and mRNA expression dysregulation on survival of all TCGA BLCA patients (n=413). Plot was obtained from cBioPortal. (B) EZH2 activity is affected by PAK4 inhibition. IPA upstream regulator analysis predicted the potential inhibition of EZH2 activity upon PAK4 inhibition. The derepression of genes (CYP1B1, CXCL8, C3, BIRC3, SLC1A3, PCDHB5, NFKBIA, MPZL2, LCN2 and DKK1 regulated by EZH2) was observed upon PAK4 inhibition. Green and red colored nodes indicate genes that are down-regulated and up-regulated upon PAK4 inhibitor treatment, respectively.

**Supplementary Figure 3:** (A) The predicted binding sites of miR-122 and miR-193 in the 3'-UTR of PAK4. (B) Western blot analysis showing PAK4 in lysates of VM-CUB1 and RT-112 cells transiently overexpressing PAK4 and treated with NT pre-miR or pre-miR-27a, -122, -128, -193, or -217. β-Actin was used as a loading control. (C) Colony formation efficiency was low for cells treated with pre-miR-122 or 193, as compared with NT miR.

**Supplementary Figure 4:** IPA (Ingenuity Pathway Analysis) led to identification of enriched canonical pathways associated with genes differentially expressed on (A) PAK4 inhibitor and (B) PAK4 siRNA treatment. Top 10 pathways found by IPA's Core analysis are listed, and canonical pathways are arranged based on significance level (-log [P-value]).

**Supplementary Figure 5:** Protein interaction network of significantly differentially expressed genes. (A) The network represents genes significantly up/down-regulated in VMCUB1 bladder cancer cell line treated with inhibitor and/or siRNA against PAK4. Protein interactions with a minimum confidence threshold of 0.5 from STRING were analyzed using Cytoscape. Primary and secondary interactions of PAK4 are highlighted with different node sizes. Hubs identified within network are highlighted.

Chandrashekar et al., Supplementary Figure 1

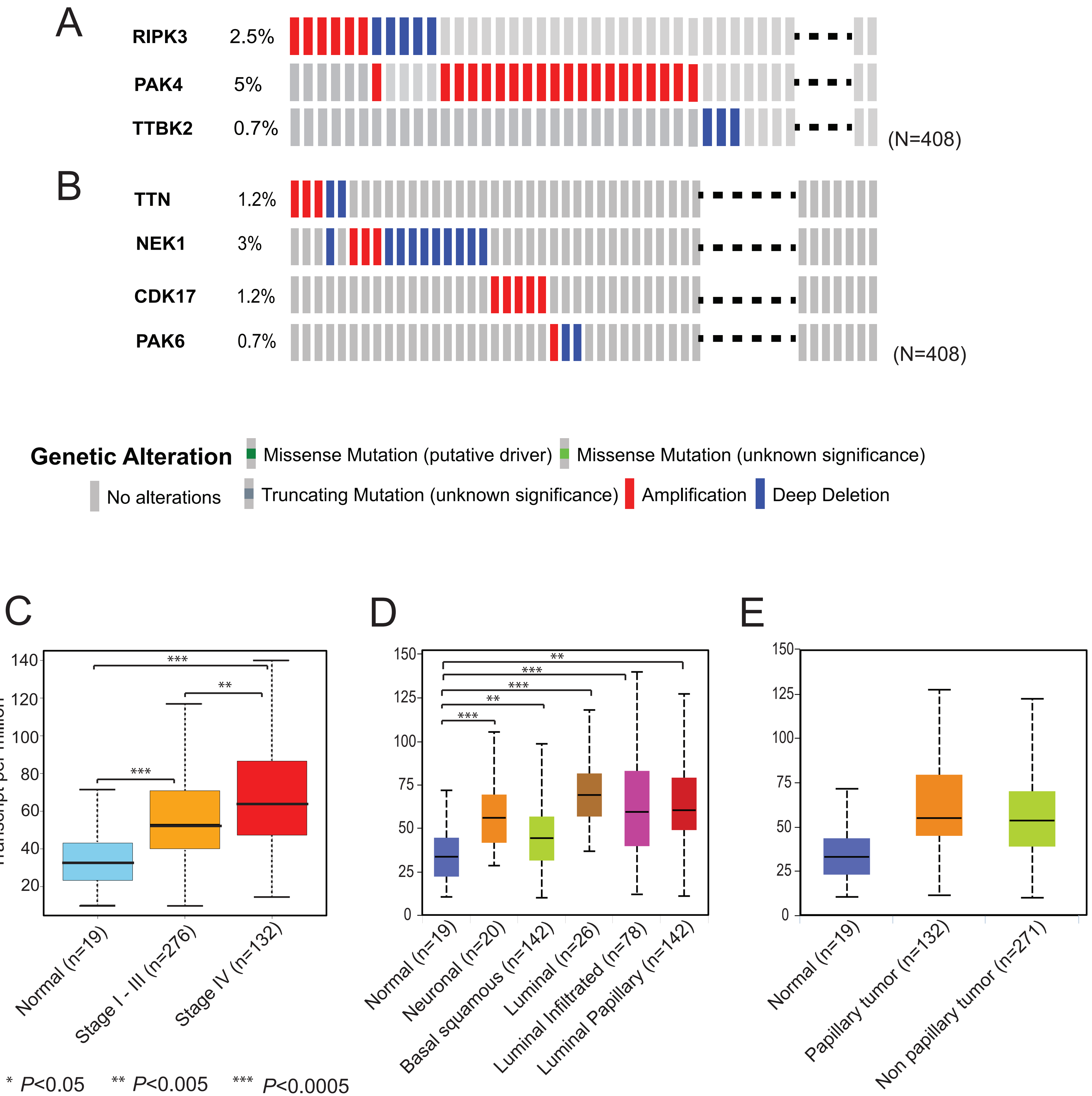

**Chandrashekar et al., Supplementary Figure 2**

A

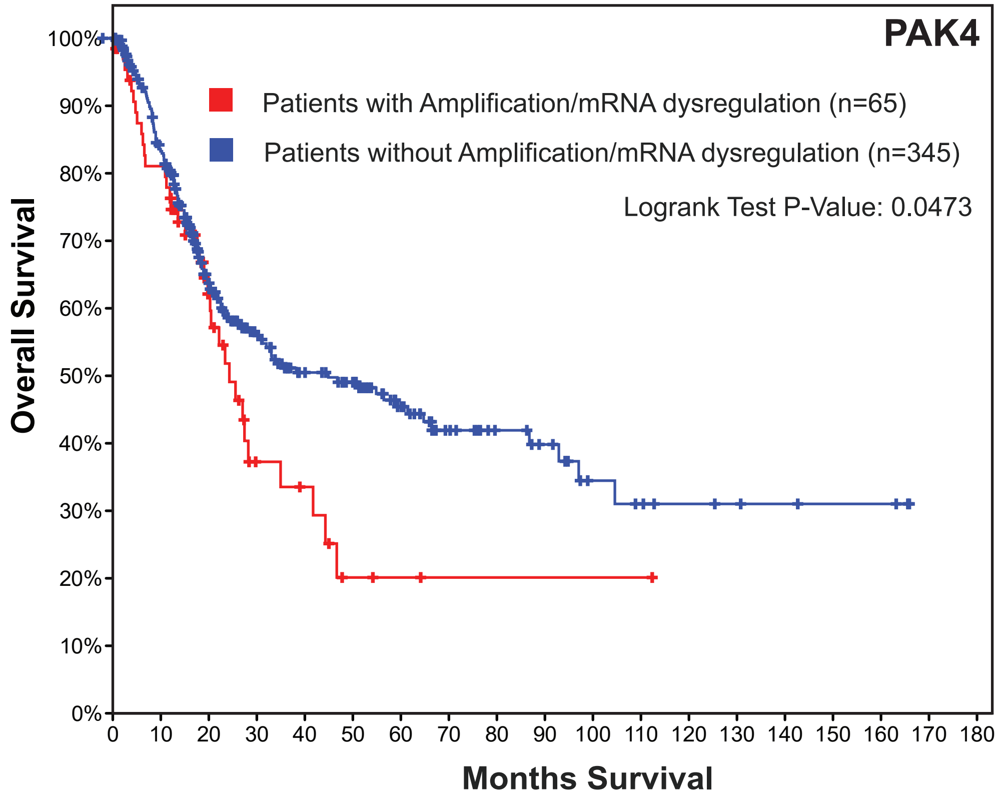


B

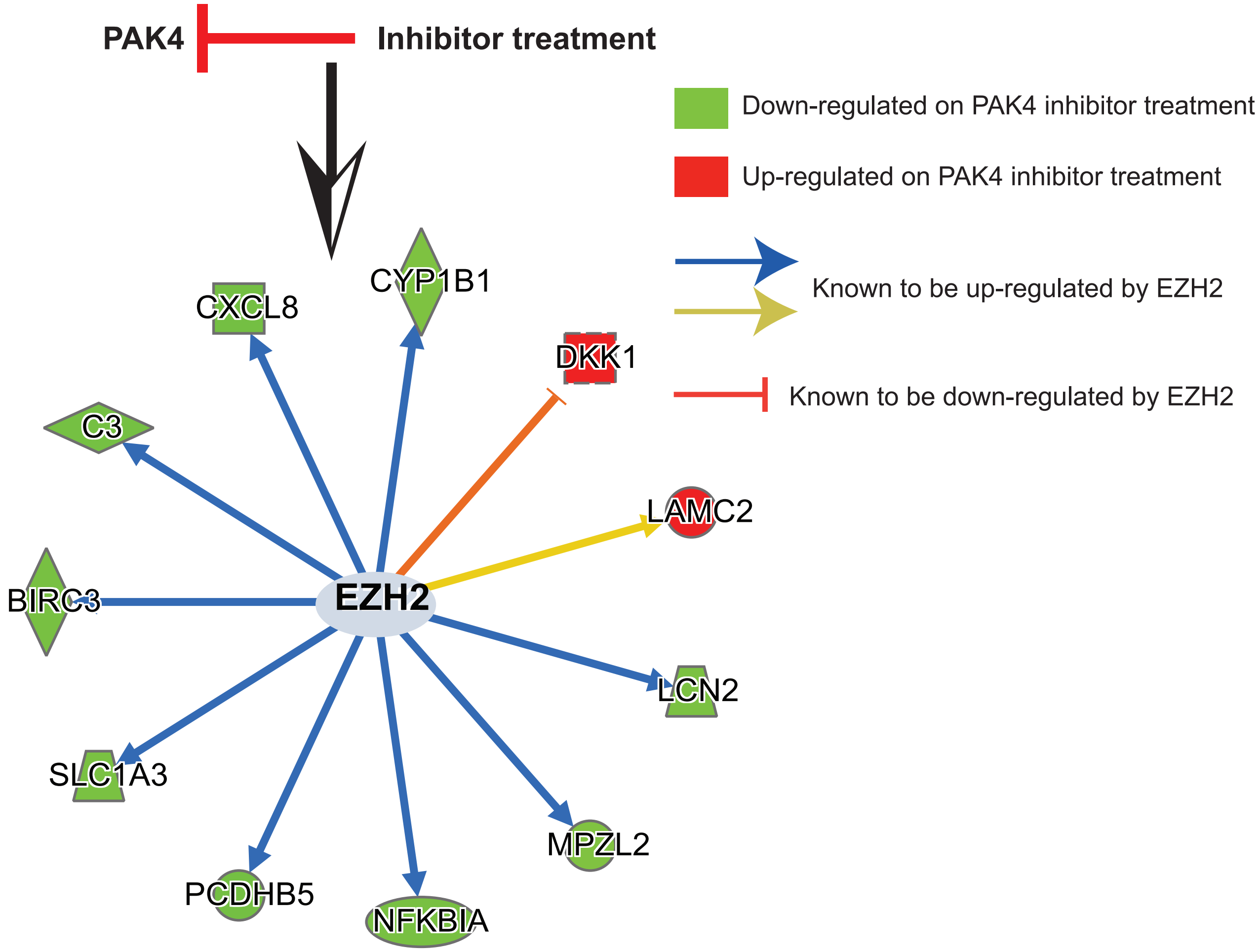

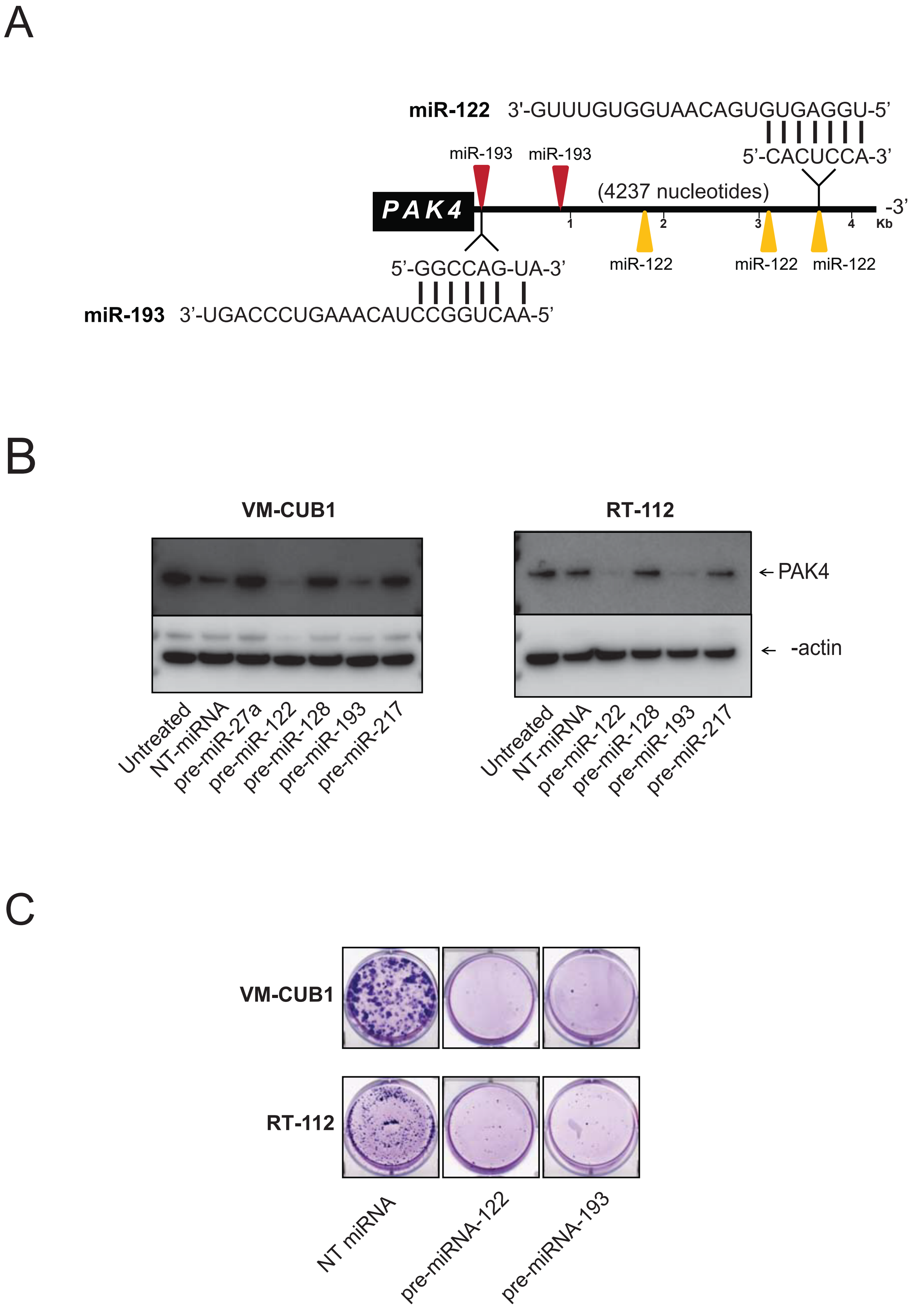
A
miR-122 3'-GUUUGUGGUAACAGUGUGAGGU-5'
5'-CACUCCA-3'
miR-193
miR-193
(4237 nucleotides)
PAK4
-3'
1
2
3
4
Kb
miR-122
miR-122
miR-122
5'-GGCCAG-UA-3'
miR-193 3'-UGACCCUGAAACAUCCGGUCAA-5'
B
VM-CUB1
RT-112
PAK4
-actin
Untreated
NT-miRNA
pre-miR-27a
pre-miR-122
pre-miR-128
pre-miR-193
pre-miR-217
Untreated
NT-miRNA
pre-miR-122
pre-miR-128
pre-miR-193
pre-miR-217
C
VM-CUB1
RT-112
NT miRNA
pre-miRNA-122
pre-miRNA-193

**Chandrashekar et al., Supplementary Figure 4**

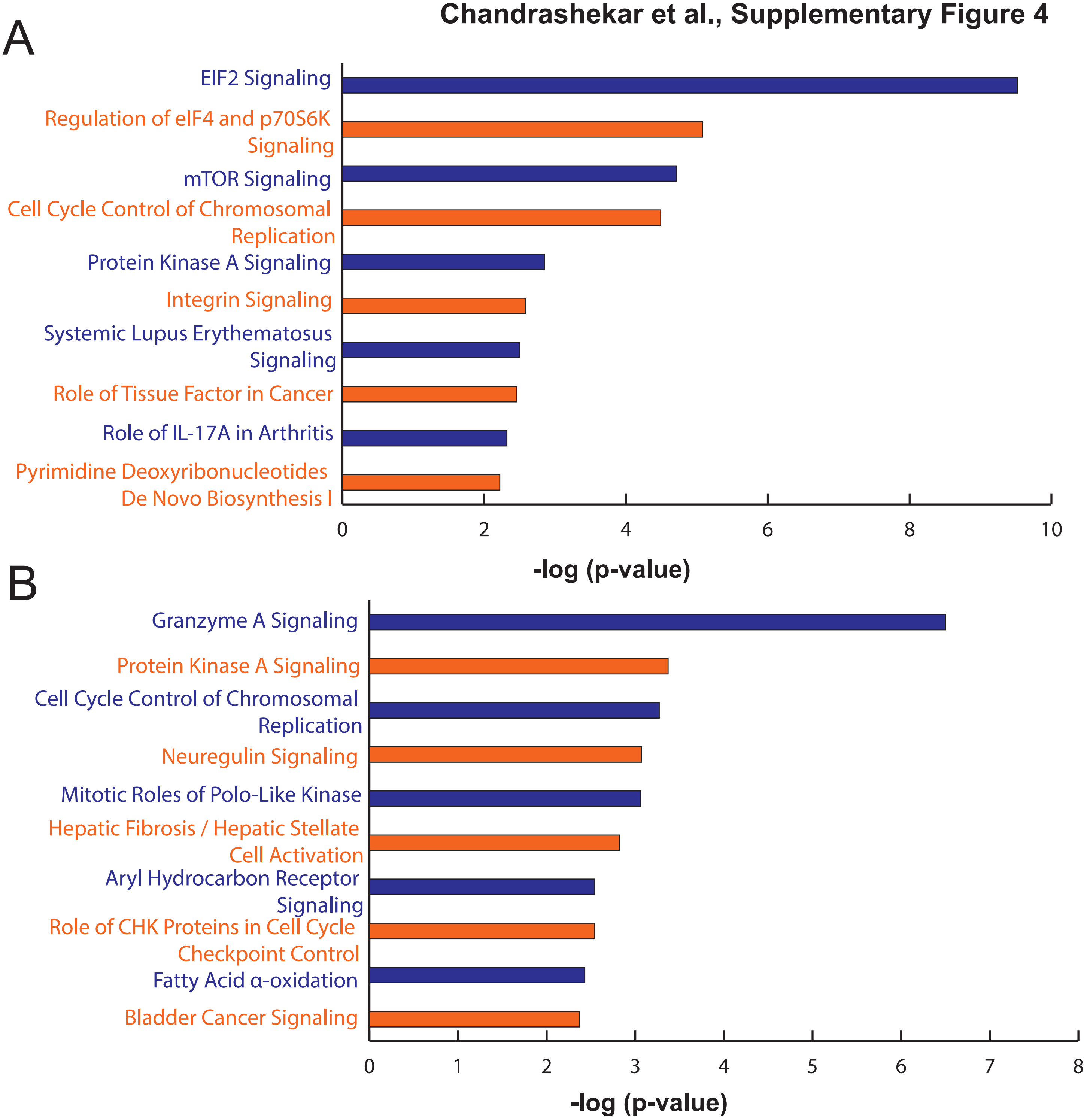

Chandrashekar et al., Supplementary Figure 5

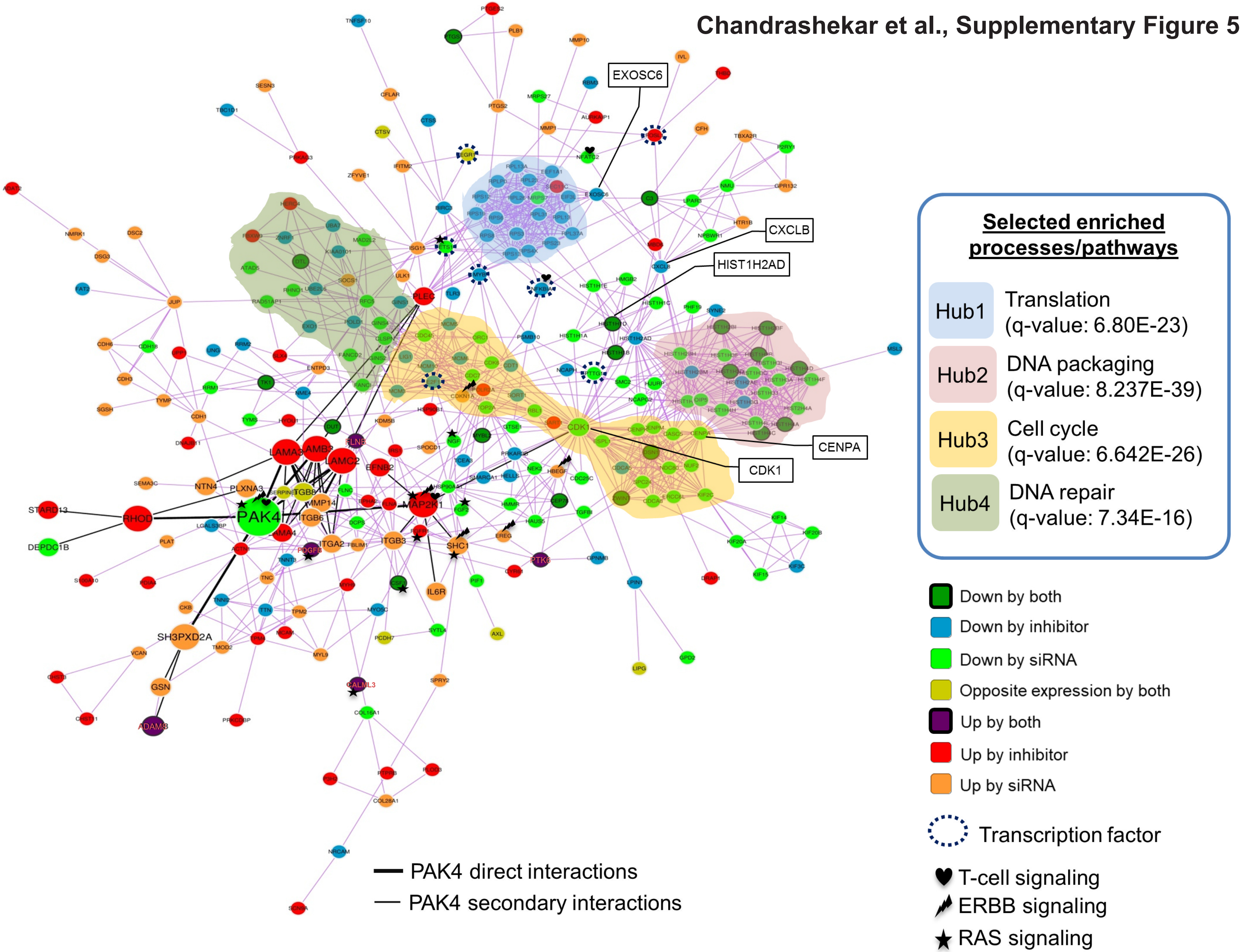
