## Supplementary tables for "Therapeutically actionable PAK4 is amplified, overexpressed and involved in bladder cancer progression"

### Shared primary authors

*Shared senior authors

Correspondence to:
Guru Sonpavde, M.D.
Bladder Cancer Director, GU Oncology division,
Dana-Farber Cancer Institute, D1230
Boston, MA, USA 02215


And
Sooryanarayana Varambally, Ph.D., Molecular and Cellular Pathology,
Department of Pathology,
Wallace Tumor Institute, Room # 430,

University of Alabama at Birmingham,
Birmingham, AL 35233, USA


**Running Title:** PAK4 amplification induces bladder cancer progression

**Disclosure of Potential Conflicts of Interest:**
Guru Sonpavde, MD was a consultant for BMS, Exelixis, Bayer, Sanofi, Pfizer, Novartis, Eisai, Janssen, Amgen, AstraZeneca, Merck, Genentech, Astellas/Agensys; Research support to institution from Bayer, Amgen, Boehringer-Ingelheim, Merck, Sanofi, Pfizer; Author for Up-to-date; Speaker for Clinical Care Options, Physicians Education Resource (PER), Research to Practice (RTP), Onclive
Christopher Willey, M.D., Ph.D., was a consultant for Varian Medical Systems and LifeNet Health, Inc.
Akhilesh Bajpai, M.Sc., and Sravanthi Duvuluri, M.Sc., received financial support from Shodhaka Life Sciences Pvt. Ltd.
Kshitish Acharya, Ph.D. is the founder and director of Shodhaka LS Pvt. Ltd
George J Netto, M.D. served as a consultant to Genentec.
Eddy S. Yang, M.D., Ph.D. was a consultant for Astrazeneca; Research support to institution from Astrazeneca, Eli Lilly, Novartis

**Supplementary Table 1:** Histological and molecular subtype of TCGA BLCA samples with PAK4 amplification. Molecular subtype information was obtained from Robertson et al.(1) Tumor histology was obtained from the clinical data for TCGA patients.

| Patient ID | Molecular subtype | Histology |
|---|---|---|
| TCGA-4Z-AA80 | Luminal_papillary | Papillary |
| TCGA-4Z-AA84 | Basal_squamous | Papillary |
| TCGA-4Z-AA87 | Luminal_infiltrated | Non-papillary |
| TCGA-5N-A9KI | Luminal_infiltrated | Non-papillary |
| TCGA-BT-A2LA | Neuronal | Non-papillary |
| TCGA-BT-A2LD | Basal_squamous | Non-papillary |
| TCGA-BT-A20O | Basal_squamous | Non-papillary |
| TCGA-BT-A20R | Luminal_infiltrated | Non-papillary |
| TCGA-CF-A1HS | Basal_squamous | Non-papillary |
| TCGA-DK-A1A5 | Luminal_infiltrated | Non-papillary |
| TCGA-DK-A1A7 | Luminal | Non-papillary |
| TCGA-DK-A1AD | Luminal | Non-papillary |
| TCGA-DK-A2HX | Luminal_infiltrated | Non-papillary |
| TCGA-FD-A3SJ | Luminal | Non-papillary |
| TCGA-FD-A3SR | Luminal_infiltrated | Non-papillary |
| TCGA-FD-A6TG | Luminal_infiltrated | Non-papillary |
| TCGA-FD-A43P | Luminal_infiltrated | Non-papillary |
| TCGA-GU-A767 | Luminal | Papillary |
| TCGA-SY-A9G5 | Basal_squamous | Papillary |
| TCGA-ZF-AA58 | Basal_squamous | Non-papillary |

**Supplementary Table 2:** Differentially expressed protein coding genes on PAK4 inhibitor treatment or PAK4 knockdown in VM-CUB1 cell line.

| PAK4 inhibitor vs DMSO | | | | PAK4 siRNA vs NT siRNA | | |
|---|---|---|---|---|---|---|
| **Gene symbol** | **Absolute FC** | **pval** | | **Gene symbol** | **Absolute FC** | **pval** |
| TMEM59L | Inf | 1.13E-02 | | MMP10 | Inf | 5.42E-04 |
| PAMR1 | 18.25 | 4.20E-03 | | COL28A1 | Inf | 3.85E-02 |
| AHRR | 12.97 | 1.64E-06 | | HES2 | 117.25 | 2.22E-05 |
| SEMA7A | 12.49 | 7.06E-03 | | SPRR1B | 96.83 | 2.46E-07 |
| PIPOX | 9.93 | 2.42E-02 | | MMP1 | 66.50 | 7.85E-04 |
| PRKAG3 | 9.28 | 3.28E-02 | | KRT13 | 61.25 | 1.23E-03 |
| THAP8 | 8.64 | 1.88E-02 | | TAGLN | 58.73 | 3.12E-08 |
| USP27X | 7.11 | 2.50E-02 | | SPRR3 | 56.00 | 1.98E-03 |
| PDGFB | 5.82 | 6.47E-04 | | KLHDC7B | 41.13 | 5.91E-04 |
| PP2D1 | 5.76 | 2.69E-02 | | ITGB3 | 41.13 | 5.91E-04 |
| RTEL1-TNFRSF6I | 5.34 | 1.96E-02 | | C8orf4 | 38.50 | 1.13E-02 |
| CABYR | 4.95 | 9.71E-03 | | C12orf54 | 36.75 | 1.37E-02 |
| SERPINI1 | 4.45 | 7.94E-05 | | WFDC3 | 35.00 | 1.66E-02 |
| FOSL1 | 4.29 | 1.08E-03 | | ENTPD3 | 31.31 | 1.17E-05 |
| HTR7 | 3.84 | 2.64E-02 | | HTR1B | 29.75 | 3.02E-02 |
| MTRNR2L12 | 3.81 | 1.98E-04 | | MYL9 | 26.89 | 1.07E-06 |
| CST6 | 3.61 | 3.39E-04 | | KPNA7 | 21.88 | 1.36E-02 |
| AKAP12 | 3.50 | 5.09E-03 | | SCEL | 21.00 | 4.82E-03 |
| CYSRT1 | 3.29 | 3.38E-03 | | SH3TC1 | 20.62 | 2.04E-05 |
| FBXW9 | 3.26 | 2.93E-02 | | SDCBP2 | 20.42 | 5.52E-03 |
| FGFR1 | 3.24 | 2.53E-03 | | CDH6 | 20.12 | 1.90E-02 |
| ADAM8 | 3.23 | 3.41E-02 | | XG | 19.25 | 2.25E-02 |
| ZNF57 | 3.17 | 4.66E-02 | | GBP6 | 18.59 | 8.08E-05 |
| FST | 3.11 | 3.67E-05 | | CLIC3 | 18.59 | 4.66E-04 |
| CMTM7 | 3.08 | 2.43E-02 | | PCDH7 | 17.40 | 9.55E-05 |
| MTRNR2L1 | 2.98 | 2.60E-05 | | TBXA2R | 16.63 | 3.81E-02 |
| GCAT | 2.94 | 2.64E-02 | | IVL | 15.75 | 4.56E-02 |
| CTSV | 2.91 | 3.47E-02 | | KRT17 | 15.74 | 1.30E-05 |
| SERPINE1 | 2.84 | 4.84E-04 | | COX6B2 | 14.44 | 1.18E-02 |
| BNC1 | 2.84 | 6.92E-03 | | HAS3 | 14.05 | 1.23E-04 |
| ADAT2 | 2.82 | 3.23E-02 | | VCAN | 13.77 | 3.20E-05 |
| KRT6A | 2.76 | 9.12E-05 | | CFH | 12.87 | 2.08E-04 |
| PRKCDBP | 2.73 | 4.55E-02 | | KRT6B | 12.03 | 1.07E-03 |
| FAM83C | 2.72 | 7.95E-03 | | GJB6 | 11.40 | 3.78E-04 |
| POLR2A | 2.70 | 9.59E-05 | | AIM2 | 11.37 | 3.89E-03 |
| ANO1 | 2.67 | 9.46E-04 | | ORAI3 | 11.37 | 1.24E-02 |
| DKK1 | 2.67 | 4.24E-03 | | AHNAK2 | 11.32 | 1.06E-04 |
| FLNA | 2.64 | 8.31E-05 | | S100A9 | 11.20 | 4.21E-03 |
| LGMN | 2.63 | 2.63E-03 | | PVRL4 | 10.85 | 4.92E-03 |
| ADAMTSL4 | 2.62 | 2.95E-02 | | TSPAN1 | 10.24 | 2.13E-04 |
| STARD13 | 2.60 | 4.09E-02 | | LYPD3 | 10.09 | 2.47E-03 |

| | | | | | | |
|---|---|---|---|---|---|---|
| ADAMTS1 | 2.60 | 4.63E-03 | | DHRS9 | 9.92 | 2.19E-02 |
| SCHIP1 | 2.54 | 4.96E-02 | | SEMA7A | 9.83 | 2.30E-04 |
| SCN9A | 2.52 | 3.69E-02 | | PTK6 | 9.73 | 5.53E-04 |
| LAMC2 | 2.50 | 1.99E-04 | | ZG16B | 9.45 | 3.77E-02 |
| FBXL6 | 2.49 | 1.83E-02 | | ST3GAL5 | 9.41 | 1.51E-02 |
| SLX4 | 2.48 | 4.44E-02 | | IQSEC2 | 9.33 | 2.77E-02 |
| SEMA3B | 2.47 | 3.96E-03 | | ABCA12 | 9.08 | 4.69E-03 |
| ABTB2 | 2.46 | 3.16E-02 | | RGS2 | 9.00 | 9.15E-04 |
| PTK6 | 2.44 | 2.54E-03 | | SOCS1 | 8.75 | 4.92E-02 |
| BCL6 | 2.39 | 1.42E-02 | | ARID3A | 8.59 | 1.20E-02 |
| AGR2 | 2.35 | 2.00E-02 | | FUT3 | 8.53 | 3.32E-03 |
| PTPRB | 2.33 | 3.93E-02 | | SMIM14 | 8.53 | 2.25E-02 |
| LZTR1 | 2.33 | 7.97E-03 | | HAL | 8.49 | 4.50E-03 |
| CYR61 | 2.33 | 8.94E-03 | | GPR157 | 8.48 | 2.21E-03 |
| AXL | 2.33 | 1.29E-03 | | EGR1 | 8.09 | 2.77E-02 |
| FAM43A | 2.31 | 4.72E-02 | | GPR132 | 7.97 | 2.38E-02 |
| UPP1 | 2.22 | 1.53E-03 | | ABCA7 | 7.93 | 1.45E-03 |
| IDNK | 2.20 | 4.88E-02 | | ULBP2 | 7.67 | 1.46E-02 |
| FADS3 | 2.20 | 1.61E-03 | | MXD4 | 7.50 | 3.16E-03 |
| MYEOV | 2.20 | 9.08E-03 | | FXYD3 | 7.50 | 1.23E-03 |
| TM7SF2 | 2.20 | 1.95E-02 | | WIPI1 | 7.39 | 5.74E-03 |
| NETO2 | 2.19 | 2.21E-02 | | FAM214B | 7.37 | 1.54E-02 |
| PHLDA1 | 2.18 | 3.20E-03 | | TYMP | 7.32 | 2.24E-03 |
| PTPRH | 2.17 | 3.39E-02 | | MAST4 | 7.26 | 1.38E-03 |
| ADTRP | 2.16 | 3.49E-02 | | STEAP4 | 7.16 | 5.38E-03 |
| THBD | 2.15 | 3.93E-02 | | ITGB8 | 7.13 | 6.81E-03 |
| DPAGT1 | 2.14 | 1.55E-02 | | RNF223 | 6.93 | 9.10E-03 |
| FLNB | 2.11 | 2.16E-03 | | KRT6C | 6.82 | 3.39E-03 |
| KRT80 | 2.08 | 1.12E-02 | | SLC25A23 | 6.60 | 4.51E-03 |
| SEC11C | 2.08 | 1.74E-02 | | EREG | 6.60 | 1.90E-03 |
| CYP4F11 | 2.07 | 1.53E-02 | | NTN4 | 6.49 | 5.04E-03 |
| EFNB2 | 2.07 | 7.40E-03 | | PGPEP1 | 6.48 | 1.57E-02 |
| CRELD2 | 2.07 | 3.94E-03 | | HEG1 | 6.47 | 2.35E-03 |
| F3 | 2.05 | 3.25E-03 | | MVP | 6.42 | 2.79E-03 |
| MBD6 | 2.05 | 3.18E-02 | | NMRK1 | 6.39 | 7.92E-03 |
| CALML3 | 2.04 | 4.32E-02 | | GIPR | 6.38 | 2.79E-02 |
| PLEC | 2.04 | 3.66E-03 | | HAS2 | 6.28 | 2.24E-02 |
| PTRF | 2.00 | 7.64E-03 | | PRSS22 | 6.04 | 6.02E-03 |
| P3H2 | 1.99 | 4.90E-02 | | HSPB1 | 6.00 | 3.48E-03 |
| TINAGL1 | 1.98 | 5.40E-03 | | PTGS2 | 5.95 | 1.37E-02 |
| LAMA4 | 1.97 | 4.11E-02 | | CALML3 | 5.95 | 5.09E-03 |
| C11orf68 | 1.97 | 3.17E-02 | | TMCC3 | 5.93 | 2.61E-02 |
| MCAM | 1.95 | 1.54E-02 | | KRT16 | 5.92 | 1.65E-02 |
| BCAR3 | 1.95 | 3.28E-02 | | IFITM2 | 5.87 | 2.94E-02 |
| LTBP4 | 1.95 | 4.38E-02 | | TMC6 | 5.82 | 1.12E-02 |
| TUFT1 | 1.94 | 1.73E-02 | | C6orf223 | 5.79 | 1.60E-02 |
| FHL2 | 1.94 | 2.10E-02 | | RBMS3 | 5.75 | 2.43E-02 |
| PLOD3 | 1.94 | 2.34E-02 | | TMOD2 | 5.67 | 2.57E-02 |
| COTL1 | 1.93 | 1.72E-02 | | MMP14 | 5.67 | 3.76E-03 |
| ABCA7 | 1.93 | 4.02E-02 | | ULK1 | 5.64 | 1.43E-02 |

| | | | | | | |
|---|---|---|---|---|---|---|
| ANXA3 | 1.93 | 1.37E-02 | | TP53INP2 | 5.61 | 1.55E-02 |
| KRT7 | 1.92 | 8.04E-03 | | IKZF2 | 5.54 | 1.54E-02 |
| CHST3 | 1.91 | 2.76E-02 | | SEMA4A | 5.52 | 1.72E-02 |
| S100A10 | 1.90 | 1.10E-02 | | CDH1 | 5.43 | 4.73E-03 |
| AMIGO2 | 1.89 | 1.26E-02 | | GBP1 | 5.40 | 8.40E-03 |
| DNAJB11 | 1.86 | 1.26E-02 | | GRAMD2 | 5.33 | 3.27E-02 |
| C10orf54 | 1.86 | 1.71E-02 | | PDCD1LG2 | 5.33 | 1.47E-02 |
| RHOD | 1.84 | 2.03E-02 | | TMC7 | 5.31 | 1.24E-02 |
| EPHA2 | 1.84 | 1.79E-02 | | SLC6A9 | 5.25 | 1.13E-02 |
| ASAP1 | 1.83 | 2.51E-02 | | SEMA3C | 5.23 | 5.60E-03 |
| HSP90B1 | 1.83 | 1.22E-02 | | CDKN1A | 5.15 | 6.78E-03 |
| DRAP1 | 1.83 | 2.38E-02 | | TACSTD2 | 5.11 | 6.16E-03 |
| AURKAIP1 | 1.82 | 4.26E-02 | | SERPINI1 | 5.07 | 2.09E-02 |
| CHST11 | 1.81 | 3.39E-02 | | LIPG | 5.06 | 4.79E-02 |
| HYOU1 | 1.80 | 1.54E-02 | | NLRP1 | 5.02 | 1.66E-02 |
| SART1 | 1.79 | 4.37E-02 | | CDH3 | 5.00 | 6.91E-03 |
| ARFIP2 | 1.76 | 4.67E-02 | | ABCG1 | 4.96 | 3.72E-02 |
| MAP2K1 | 1.76 | 3.92E-02 | | GPR153 | 4.94 | 1.18E-02 |
| PTGES2 | 1.75 | 4.87E-02 | | ADM2 | 4.92 | 4.44E-02 |
| G6PD | 1.74 | 4.59E-02 | | AREG | 4.91 | 7.75E-03 |
| EMC7 | 1.73 | 4.66E-02 | | GJB2 | 4.90 | 8.33E-03 |
| PHLDB2 | 1.72 | 3.07E-02 | | GSN | 4.88 | 1.73E-02 |
| ASAP2 | 1.71 | 4.24E-02 | | ADAM8 | 4.87 | 1.07E-02 |
| TPM4 | 1.71 | 3.60E-02 | | PCDH1 | 4.86 | 1.03E-02 |
| ACTN1 | 1.70 | 3.50E-02 | | PLAT | 4.85 | 3.73E-02 |
| MYH9 | 1.69 | 3.01E-02 | | PLB1 | 4.85 | 3.73E-02 |
| IRS1 | 1.68 | 4.77E-02 | | ITGB6 | 4.84 | 7.97E-03 |
| ZMIZ1 | 1.68 | 4.96E-02 | | SERINC5 | 4.83 | 1.14E-02 |
| LAMA3 | 1.66 | 3.75E-02 | | FUCA1 | 4.80 | 1.47E-02 |
| HERC4 | 1.65 | 4.98E-02 | | S100A14 | 4.80 | 1.61E-02 |
| PDIA4 | 1.64 | 4.14E-02 | | TM7SF2 | 4.79 | 2.71E-02 |
| LAMB3 | 1.63 | 4.16E-02 | | TRPM4 | 4.65 | 1.70E-02 |
| RPL23 | -1.62 | 4.79E-02 | | MACC1 | 4.63 | 2.22E-02 |
| SORL1 | -1.63 | 4.70E-02 | | PDGFB | 4.60 | 1.18E-02 |
| RPS19 | -1.65 | 4.18E-02 | | KCTD11 | 4.58 | 1.60E-02 |
| GLTSCR2 | -1.65 | 4.98E-02 | | TNC | 4.54 | 1.15E-02 |
| EIF3L | -1.67 | 4.53E-02 | | KRT6A | 4.52 | 1.07E-02 |
| ATAD2 | -1.68 | 4.58E-02 | | DSG3 | 4.50 | 1.34E-02 |
| NFKBIA | -1.68 | 4.50E-02 | | DUSP4 | 4.48 | 1.45E-02 |
| ALDH3A1 | -1.72 | 2.56E-02 | | ZFYVE1 | 4.45 | 2.34E-02 |
| RPL13 | -1.73 | 2.64E-02 | | MAFK | 4.40 | 1.52E-02 |
| LAMP3 | -1.73 | 3.84E-02 | | JUP | 4.40 | 1.25E-02 |
| RPS6 | -1.73 | 2.38E-02 | | ITGA2 | 4.35 | 1.31E-02 |
| RPL31 | -1.73 | 2.76E-02 | | S100A6 | 4.31 | 1.38E-02 |
| SREBF1 | -1.74 | 3.08E-02 | | LRRN1 | 4.29 | 3.00E-02 |
| ID1 | -1.77 | 2.33E-02 | | TMEM150A | 4.26 | 4.39E-02 |
| RPLP0 | -1.78 | 2.00E-02 | | ADGRF4 | 4.24 | 4.89E-02 |
| UBE2L6 | -1.78 | 4.15E-02 | | KDM7A | 4.23 | 2.57E-02 |
| RIMS2 | -1.79 | 4.59E-02 | | KYNU | 4.23 | 2.16E-02 |
| PLTP | -1.79 | 3.45E-02 | | YPEL5 | 4.17 | 2.29E-02 |

| | | | | | | |
|---|---|---|---|---|---|---|
| RPL37A | -1.80 | 1.65E-02 | | ARL8A | 4.15 | 3.44E-02 |
| TK1 | -1.80 | 4.98E-02 | | ISG15 | 4.12 | 2.15E-02 |
| LGALS3BP | -1.80 | 4.03E-02 | | BHLHE40 | 4.08 | 2.49E-02 |
| ADGRF1 | -1.82 | 1.86E-02 | | PPM1K | 4.07 | 4.62E-02 |
| ELF3 | -1.82 | 1.65E-02 | | SGSH | 4.06 | 4.19E-02 |
| PELI2 | -1.83 | 4.62E-02 | | INHBA | 4.03 | 3.35E-02 |
| RPL26 | -1.83 | 2.43E-02 | | IFFO2 | 3.96 | 1.99E-02 |
| HIST1H3B | -1.83 | 2.99E-02 | | SPOCD1 | 3.91 | 3.99E-02 |
| HELLS | -1.85 | 2.08E-02 | | CFLAR | 3.91 | 2.63E-02 |
| RPS23 | -1.85 | 1.43E-02 | | C7orf43 | 3.89 | 3.96E-02 |
| HIST1H2AE | -1.85 | 3.70E-02 | | SESN3 | 3.88 | 3.72E-02 |
| MANBA | -1.86 | 3.77E-02 | | DNAH5 | 3.83 | 4.74E-02 |
| FAT2 | -1.87 | 1.13E-02 | | FLNB | 3.81 | 2.27E-02 |
| RPS3 | -1.88 | 9.68E-03 | | IL1B | 3.76 | 2.62E-02 |
| CDCA7 | -1.88 | 4.29E-02 | | MPZL2 | 3.73 | 3.17E-02 |
| EEF1A1 | -1.88 | 9.36E-03 | | GALNT5 | 3.65 | 2.99E-02 |
| DDIT4 | -1.89 | 1.04E-02 | | CKB | 3.62 | 4.19E-02 |
| RPS13 | -1.89 | 1.95E-02 | | ANGPTL4 | 3.61 | 3.49E-02 |
| HIST1H2BF | -1.90 | 3.71E-02 | | SHC1 | 3.60 | 2.90E-02 |
| RPS12 | -1.90 | 1.11E-02 | | C3orf52 | 3.56 | 4.00E-02 |
| HIST1H2BI | -1.91 | 4.59E-02 | | DUSP6 | 3.56 | 3.81E-02 |
| RAVER2 | -1.91 | 3.26E-02 | | LRP10 | 3.56 | 3.29E-02 |
| NR1H2 | -1.91 | 2.88E-02 | | KRT19 | 3.55 | 3.04E-02 |
| PTPRS | -1.92 | 1.51E-02 | | MXD1 | 3.53 | 3.78E-02 |
| SORT1 | -1.92 | 1.58E-02 | | SPRY2 | 3.52 | 4.87E-02 |
| SLC23A2 | -1.92 | 2.73E-02 | | PLXNA3 | 3.49 | 4.55E-02 |
| HIST1H4C | -1.93 | 1.42E-02 | | KRT5 | 3.48 | 3.31E-02 |
| ZWINT | -1.93 | 3.55E-02 | | FBLIM1 | 3.47 | 3.67E-02 |
| IKZF2 | -1.94 | 1.35E-02 | | IL6R | 3.47 | 3.96E-02 |
| IFI44 | -1.94 | 3.82E-02 | | CD24 | 3.46 | 3.47E-02 |
| RBM3 | -1.95 | 9.83E-03 | | KIAA1161 | 3.44 | 4.22E-02 |
| KIAA0101 | -1.95 | 3.89E-02 | | MSMO1 | 3.42 | 3.72E-02 |
| RPL13A | -1.95 | 6.55E-03 | | TNFAIP8 | 3.41 | 4.60E-02 |
| HIST1H1D | -1.96 | 1.95E-02 | | HBEGF | 3.37 | 4.21E-02 |
| RPS4X | -1.96 | 6.97E-03 | | EPPK1 | 3.36 | 4.48E-02 |
| RPS8 | -1.96 | 5.95E-03 | | ALDH1L2 | 3.36 | 4.38E-02 |
| CEP78 | -1.97 | 2.33E-02 | | PERP | 3.32 | 4.01E-02 |
| GINS1 | -1.98 | 3.89E-02 | | SH3PXD2A | 3.31 | 4.24E-02 |
| ADD3 | -1.99 | 1.10E-02 | | KDM5B | 3.30 | 4.43E-02 |
| ARRDC3 | -1.99 | 8.89E-03 | | DDIT4 | 3.29 | 4.58E-02 |
| HIST1H3D | -1.99 | 2.64E-02 | | ERRFI1 | 3.28 | 4.41E-02 |
| BMF | -2.00 | 1.48E-02 | | DSC2 | 3.27 | 4.71E-02 |
| LCN2 | -2.01 | 4.15E-03 | | TPM2 | 3.26 | 4.51E-02 |
| SCD | -2.02 | 4.92E-03 | | THSD4 | 3.23 | 4.95E-02 |
| EXOSC6 | -2.05 | 8.27E-03 | | CLDN1 | 3.16 | 4.93E-02 |
| UNG | -2.05 | 1.38E-02 | | MKI67 | -3.14 | 4.98E-02 |
| ATP9A | -2.07 | 7.52E-03 | | FANCI | -3.21 | 4.81E-02 |
| MPZL2 | -2.08 | 1.37E-02 | | HIST1H2BH | -3.22 | 4.93E-02 |
| PTGS1 | -2.09 | 3.55E-02 | | HIST1H4C | -3.23 | 4.73E-02 |
| MSL3 | -2.09 | 4.16E-02 | | HIST1H3F | -3.25 | 4.89E-02 |

| | | | | | | |
|---|---|---|---|---|---|---|
| ATOH8 | -2.09 | 1.94E-02 | | CLSPN | -3.25 | 4.90E-02 |
| MCM5 | -2.10 | 7.29E-03 | | MYBL2 | -3.27 | 4.55E-02 |
| COBLL1 | -2.10 | 1.48E-02 | | CDCA8 | -3.28 | 4.86E-02 |
| SYNE2 | -2.12 | 3.65E-03 | | DTL | -3.30 | 4.73E-02 |
| SLC44A3 | -2.13 | 3.97E-02 | | HMMR | -3.31 | 4.90E-02 |
| DTWD2 | -2.16 | 2.49E-02 | | TOP2A | -3.32 | 3.99E-02 |
| PIK3IP1 | -2.17 | 1.63E-02 | | RRM1 | -3.33 | 4.22E-02 |
| CDCA5 | -2.17 | 2.18E-02 | | ORC1 | -3.37 | 4.84E-02 |
| HIST1H1B | -2.17 | 4.93E-03 | | FAM83D | -3.37 | 4.30E-02 |
| STEAP4 | -2.17 | 2.03E-02 | | DEPDC1 | -3.37 | 4.44E-02 |
| KLHDC7B | -2.17 | 8.77E-03 | | ATAD5 | -3.39 | 4.45E-02 |
| TLR3 | -2.17 | 6.99E-03 | | ZWINT | -3.40 | 4.16E-02 |
| MCM10 | -2.18 | 3.62E-02 | | NEMP1 | -3.40 | 4.12E-02 |
| LIG1 | -2.19 | 2.28E-02 | | SPAG5 | -3.41 | 4.24E-02 |
| SLPI | -2.19 | 2.63E-02 | | SMC2 | -3.41 | 3.82E-02 |
| HIST1H3G | -2.21 | 1.29E-02 | | KIF20B | -3.41 | 3.88E-02 |
| SPTLC3 | -2.21 | 4.27E-02 | | TMPO | -3.44 | 3.67E-02 |
| KRT15 | -2.22 | 8.89E-03 | | WDR76 | -3.44 | 4.78E-02 |
| LY6D | -2.23 | 1.91E-03 | | MMS22L | -3.46 | 4.13E-02 |
| MRC2 | -2.24 | 1.06E-02 | | HIST1H3I | -3.47 | 4.04E-02 |
| FAM111B | -2.25 | 2.55E-02 | | CDK6 | -3.48 | 3.28E-02 |
| TBC1D1 | -2.27 | 1.38E-02 | | CEP78 | -3.49 | 3.69E-02 |
| HIST1H2AD | -2.27 | 1.90E-02 | | KIF2C | -3.49 | 3.80E-02 |
| C3 | -2.28 | 1.16E-03 | | FANCD2 | -3.49 | 4.23E-02 |
| DSN1 | -2.28 | 2.59E-02 | | HMGB2 | -3.49 | 3.39E-02 |
| MCM6 | -2.28 | 6.44E-03 | | KIAA1524 | -3.49 | 3.63E-02 |
| FAM195B | -2.31 | 2.73E-02 | | PHF19 | -3.49 | 3.99E-02 |
| DUT | -2.32 | 7.84E-03 | | ASPM | -3.50 | 3.36E-02 |
| TEF | -2.33 | 1.68E-03 | | CDK1 | -3.52 | 3.62E-02 |
| SLC1A4 | -2.35 | 2.85E-02 | | ESPL1 | -3.53 | 4.06E-02 |
| RAB26 | -2.37 | 4.18E-02 | | ERCC6L | -3.54 | 4.70E-02 |
| BIRC3 | -2.37 | 2.38E-03 | | HIST1H4A | -3.54 | 4.33E-02 |
| CDT1 | -2.38 | 1.73E-02 | | DEPDC1B | -3.55 | 4.25E-02 |
| TTC22 | -2.39 | 2.72E-02 | | RFC5 | -3.56 | 4.57E-02 |
| CTSS | -2.41 | 1.41E-02 | | HIST1H2BI | -3.58 | 3.35E-02 |
| C1R | -2.41 | 8.09E-03 | | UHMK1 | -3.61 | 2.91E-02 |
| HIST1H4D | -2.42 | 7.46E-03 | | CDCA3 | -3.61 | 4.40E-02 |
| PSMB10 | -2.42 | 1.13E-02 | | HIST1H4H | -3.62 | 3.12E-02 |
| GPR85 | -2.42 | 5.40E-03 | | EIF2B1 | -3.62 | 3.74E-02 |
| SLC2A10 | -2.44 | 4.60E-02 | | KIF14 | -3.63 | 3.14E-02 |
| TP53INP1 | -2.46 | 1.14E-03 | | TK1 | -3.63 | 3.05E-02 |
| SMARCA1 | -2.48 | 9.84E-03 | | HIST1H2BF | -3.67 | 3.23E-02 |
| DLK2 | -2.48 | 8.04E-03 | | HSP90AA1 | -3.67 | 2.62E-02 |
| DTL | -2.48 | 3.83E-03 | | HAUS5 | -3.68 | 4.36E-02 |
| EXO1 | -2.51 | 1.96E-02 | | HIST1H3B | -3.72 | 2.69E-02 |
| SSPN | -2.53 | 9.94E-03 | | CASC5 | -3.73 | 2.70E-02 |
| APOL6 | -2.54 | 2.67E-04 | | CDC7 | -3.76 | 3.36E-02 |
| MCM3 | -2.54 | 7.81E-04 | | HMGN2 | -3.77 | 3.33E-02 |
| AKNA | -2.54 | 1.19E-02 | | HIST1H1E | -3.78 | 2.49E-02 |
| POLD1 | -2.55 | 3.75E-03 | | MRPS7 | -3.78 | 2.94E-02 |

| | | | | | | |
|---|---|---|---|---|---|---|
| SESTD1 | -2.55 | 4.11E-02 | | ZGRF1 | -3.79 | 4.67E-02 |
| TNNI2 | -2.58 | 7.19E-03 | | DUT | -3.80 | 2.73E-02 |
| HIST1H4A | -2.60 | 1.05E-02 | | RAD51AP1 | -3.81 | 4.34E-02 |
| ZNRF1 | -2.60 | 2.37E-02 | | HIST1H1C | -3.82 | 2.34E-02 |
| CSMD1 | -2.68 | 4.77E-02 | | ETS1 | -3.83 | 2.46E-02 |
| NEBL | -2.68 | 2.33E-03 | | GTSE1 | -3.83 | 3.13E-02 |
| SMO | -2.73 | 1.58E-02 | | HIST1H4F | -3.84 | 3.44E-02 |
| AQP3 | -2.73 | 5.43E-03 | | DHX40 | -3.85 | 3.30E-02 |
| TGM1 | -2.78 | 3.83E-03 | | MRPS27 | -3.87 | 2.88E-02 |
| CXXC4 | -2.79 | 2.35E-02 | | HIST1H2AL | -3.88 | 3.24E-02 |
| GALM | -2.80 | 6.31E-03 | | CD83 | -3.90 | 3.31E-02 |
| TTN | -2.82 | 3.98E-02 | | HIST1H3A | -3.91 | 3.00E-02 |
| TSNARE1 | -2.85 | 4.89E-02 | | E2F8 | -3.92 | 4.45E-02 |
| PCDHB5 | -2.86 | 4.47E-02 | | GINS2 | -3.92 | 3.07E-02 |
| TNFRSF9 | -2.95 | 2.50E-03 | | PSRC1 | -3.94 | 3.19E-02 |
| HIST1H2AL | -2.97 | 2.53E-02 | | OLFML2A | -3.94 | 4.94E-02 |
| C11orf70 | -2.97 | 3.89E-02 | | CENPA | -3.95 | 4.15E-02 |
| SMAGP | -2.99 | 2.76E-02 | | CENPK | -3.96 | 3.04E-02 |
| CXCL8 | -3.00 | 3.93E-02 | | HIST1H3C | -3.96 | 2.25E-02 |
| LHX6 | -3.01 | 2.07E-03 | | DSN1 | -4.02 | 2.72E-02 |
| E2F1 | -3.03 | 2.37E-02 | | HSD17B11 | -4.02 | 4.20E-02 |
| RAB27B | -3.03 | 1.88E-02 | | HIST1H3D | -4.04 | 2.02E-02 |
| PITPNM3 | -3.06 | 2.04E-02 | | HIST1H1B | -4.04 | 1.80E-02 |
| ARHGEF10L | -3.10 | 9.43E-03 | | HIST1H3J | -4.06 | 2.55E-02 |
| GBP4 | -3.12 | 3.28E-03 | | CDC45 | -4.06 | 2.58E-02 |
| SSBP2 | -3.12 | 2.62E-02 | | NDC80 | -4.06 | 2.62E-02 |
| FOXQ1 | -3.12 | 2.62E-02 | | PTTG1 | -4.07 | 2.64E-02 |
| CYP1B1 | -3.16 | 8.61E-06 | | STX3 | -4.10 | 2.66E-02 |
| ZNF367 | -3.16 | 4.38E-03 | | MAD2L2 | -4.17 | 4.77E-02 |
| MYO5C | -3.19 | 6.84E-03 | | HIST1H4L | -4.20 | 1.95E-02 |
| TRIM22 | -3.23 | 9.33E-03 | | MMD | -4.24 | 2.17E-02 |
| MDK | -3.24 | 1.01E-04 | | HIST1H4D | -4.28 | 1.69E-02 |
| NPNT | -3.24 | 1.39E-04 | | CENPM | -4.30 | 3.03E-02 |
| MYBL2 | -3.26 | 2.02E-03 | | AXL | -4.33 | 1.31E-02 |
| TWSG1 | -3.26 | 1.13E-03 | | PTGS1 | -4.35 | 3.37E-02 |
| PER2 | -3.28 | 2.09E-03 | | NUSAP1 | -4.36 | 1.46E-02 |
| CSF1 | -3.30 | 9.26E-03 | | COL16A1 | -4.36 | 3.55E-02 |
| HIST1H2BM | -3.30 | 2.91E-02 | | NCAPG2 | -4.38 | 1.50E-02 |
| PKHD1L1 | -3.33 | 6.25E-03 | | NEK2 | -4.43 | 2.00E-02 |
| TNNT3 | -3.34 | 4.20E-02 | | DLGAP5 | -4.45 | 1.37E-02 |
| E2F8 | -3.39 | 2.46E-02 | | GPD2 | -4.49 | 1.36E-02 |
| LPIN1 | -3.40 | 1.77E-02 | | HIST1H1A | -4.49 | 1.48E-02 |
| RRM2 | -3.41 | 3.35E-04 | | RBL1 | -4.51 | 1.52E-02 |
| WDR76 | -3.42 | 1.08E-03 | | HJURP | -4.58 | 1.45E-02 |
| NAALADL2 | -3.49 | 3.30E-03 | | CSF1 | -4.60 | 3.93E-02 |
| TMEM200B | -3.59 | 2.45E-02 | | CHAF1B | -4.63 | 3.83E-02 |
| NCAPH | -3.64 | 3.27E-03 | | HIST1H1D | -4.66 | 1.05E-02 |
| ASS1 | -3.65 | 3.47E-05 | | LPAR3 | -4.72 | 2.59E-02 |
| PCDH7 | -3.68 | 1.72E-04 | | ALDH9A1 | -4.76 | 1.17E-02 |
| UBA7 | -3.72 | 1.45E-03 | | CTSV | -4.79 | 2.52E-02 |

| | | | | | | |
|---|---|---|---|---|---|---|
| SLC1A3 | -3.87 | 1.27E-03 | | SLC25A13 | -4.82 | 1.03E-02 |
| NME4 | -3.92 | 4.24E-02 | | KIF20A | -4.83 | 1.01E-02 |
| TNFSF10 | -4.07 | 1.81E-05 | | SPC24 | -4.91 | 1.19E-02 |
| GRAMD3 | -4.09 | 2.83E-02 | | PLCE1 | -4.94 | 2.60E-02 |
| TNFSF15 | -4.15 | 2.01E-04 | | OIP5 | -4.96 | 3.51E-02 |
| ITGB8 | -4.27 | 1.70E-03 | | HIST1H2BJ | -5.12 | 8.09E-03 |
| IQSEC2 | -4.33 | 7.98E-03 | | SERPINE1 | -5.16 | 6.19E-03 |
| SYTL5 | -4.35 | 4.60E-03 | | KIF15 | -5.18 | 9.53E-03 |
| GPNMB | -4.37 | 1.31E-04 | | FLNC | -5.18 | 3.46E-02 |
| KIF3C | -4.82 | 4.93E-02 | | NFATC2 | -5.24 | 4.00E-02 |
| EGR1 | -4.91 | 5.40E-04 | | PARPBP | -5.31 | 2.11E-02 |
| VANGL2 | -5.13 | 3.95E-04 | | TYMS | -5.41 | 8.90E-03 |
| KCNMB4 | -5.21 | 4.78E-02 | | DIO2 | -5.41 | 1.85E-02 |
| SCAMP5 | -5.21 | 4.78E-02 | | CDKN3 | -5.44 | 1.05E-02 |
| RCSD1 | -5.83 | 8.94E-03 | | METAP1D | -5.52 | 4.76E-02 |
| PRKAR2B | -6.01 | 2.54E-03 | | DCPS | -5.59 | 4.58E-02 |
| LRG1 | -6.40 | 1.70E-02 | | PIF1 | -5.61 | 2.01E-02 |
| C8orf4 | -6.42 | 2.66E-02 | | CCDC152 | -5.71 | 2.88E-02 |
| LIPG | -7.03 | 5.81E-03 | | GINS4 | -5.85 | 8.56E-03 |
| ALDH1A1 | -7.81 | 3.38E-02 | | RAB3B | -5.90 | 3.75E-03 |
| PLCL2 | -8.07 | 2.90E-02 | | NUF2 | -5.95 | 5.59E-03 |
| IL20RA | -8.85 | 1.84E-02 | | ZNF618 | -6.10 | 9.60E-03 |
| MYB | -9.02 | 4.05E-02 | | PLIN2 | -6.10 | 2.23E-02 |
| HPGD | -9.89 | 6.14E-06 | | CDH18 | -6.20 | 4.52E-02 |
| TCEA3 | -10.41 | 2.12E-02 | | P2RY1 | -6.38 | 1.02E-02 |
| NUPR1 | -11.04 | 1.75E-03 | | CNTNAP1 | -6.57 | 4.68E-02 |
| NRCAM | -12.49 | 3.38E-02 | | CDCA7 | -6.61 | 2.34E-03 |
| NEGR1 | -20.82 | 3.68E-02 | | ARHGEF40 | -6.94 | 1.03E-02 |
| | | | | FGF2 | -7.05 | 3.66E-02 |
| | | | | CDC25C | -7.06 | 1.36E-02 |
| | | | | SLC7A2 | -7.21 | 9.70E-03 |
| | | | | TGFBI | -7.54 | 8.54E-04 |
| | | | | PAK4 | -7.69 | 3.94E-03 |
| | | | | NMU | -7.79 | 1.52E-02 |
| | | | | SGCB | -7.79 | 8.85E-03 |
| | | | | DZIP1L | -7.90 | 2.38E-02 |
| | | | | MME | -8.09 | 5.69E-03 |
| | | | | SAPCD2 | -8.34 | 7.60E-03 |
| | | | | C3 | -8.41 | 8.66E-04 |
| | | | | FAM64A | -10.29 | 5.68E-03 |
| | | | | ELOVL2 | -10.86 | 1.52E-02 |
| | | | | RHNO1 | -10.90 | 1.10E-03 |
| | | | | HR | -12.00 | 9.96E-03 |
| | | | | STK33 | -12.00 | 4.77E-02 |
| | | | | NGF | -12.14 | 9.46E-03 |
| | | | | SYTL4 | -12.57 | 8.12E-03 |
| | | | | SMO | -14.29 | 9.57E-03 |
| | | | | PDK3 | -15.71 | 1.85E-02 |
| | | | | CLDN11 | -16.10 | 2.26E-05 |
| | | | | NPBWR1 | -20.00 | 4.48E-02 |

| | | | | | | |
|---|---|---|---|---|---|---|
| | | | | ADAMTS15 | -21.43 | 4.97E-03 |
| | | | | ZNF488 | -22.29 | 3.15E-02 |
| | | | | SGCE | -22.86 | 2.89E-02 |
| | | | | TSPAN6 | -23.43 | 8.44E-04 |
| | | | | TGFBR3 | -30.86 | 7.88E-04 |
| | | | | HIST2H4A | -Inf | 1.64E-03 |
| | | | | VANGL2 | -Inf | 7.36E-03 |
| | | | | PELI2 | -Inf | 1.47E-02 |
| | | | | TRMT10A | -Inf | 3.10E-02 |

**Supplementary Table 3:** Upstream regulator analysis of DEGs on PAK4 inhbition in bladder cancer cells (VM-CUB1) using Ingenuity pathway analysis (IPA)

| Upstream regulator | Predicted Activation State | Z-score | p-value | Target molecules in dataset |
|---|---|---|---|---|
| NFkB (complex) | Inhibited | -2.722 | 1.33E-02 | BIRC3,C3,CXCL8,ELF3,F3,ITGB8,LCN2,NFKBIA,TNFSF10 |
| RABL6 | Inhibited | -2 | 2.14E-02 | DUT,MCM10,MCM5,POLD1 |
| KIAA1524 | Inhibited | -2 | 1.55E-02 | CYSRT1,GPNMB,PTPRB,RHOD |
| PI3K (family) | Inhibited | -2.066 | 1.38E-03 | BIRC3,CXCL8,PTGS1,SERPINE1,SREBF1,TNFSF10 |
| IFNL1 | Inhibited | -2.39 | 4.22E-04 | APOL6,CXCL8,IFI44,LGALS3BP,TRIM22,UBE2L6 |
| EZH2 | Inhibited | -2.626 | 0.0000162 | BIRC3,C3,CXCL8,CYP1B1,DKK1,LAMC2,LCN2,MPZL2,NFKBIA,PCDHB5,SLC1A3 |
| IFNG | Inhibited | -2.228 | 4.52E-03 | APOL6,BMF,CTSS,CXCL8,DKK1,IFI44,LAMP3,NFKBIA,TGM1,TLR3,TNFSF10,TRIM22 |
| IFNA2 | Inhibited | -2.178 | 1.20E-02 | APOL6,IFI44,LGALS3BP,TNFSF10,UBE2L6 |
| ERBB2 | Inhibited | -3.018 | 2.91E-11 | CDCA5,CDCA7,CDT1,CXCL8,E2F1,E2F8,GINS1,ID1,KRT7,LAMC2,LCN2,LIG1,MCM10,MCM3,MCM5,MCM6,MYBL2,POLD1,POLR2A,RHOD,RRM2,SERPINE1,TK1,ZWINT |
| EFNA2 | Inhibited | -2 | 9.41E-03 | FOSL1,FOXQ1,KRT7,TGM1 |
| HLX | Activated | 2 | 4.76E-04 | CTSS,CXCL8,CYP1B1,EGR1 |
| NUPR1 | Activated | 4.146 | 6.99E-05 | AKAP12,CXCL8,CYR61,DSN1,E2F8,EXO1,FAM111B,FHL2,GINS1,GRAMD3,HIST1H1B,HIST1H2AL,HIST1H2BM,HIST1H3B,HIST1H4A,MCM10,PHLDA1,SERPINE1,SYNE2,UPP1,WDR76 |
| IL1RN | Activated | 2.449 | 1.12E-03 | CTSS,IFI44,LAMP3,SERPINE1,TNFSF10,TRIM22 |
| E2F6 | Activated | 2 | 1.22E-04 | E2F1,KIAA0101,LIG1,MCM3,MCM5,RRM2 |

**Supplementary Table 4:** Summary of clinical and demographic data for MIBC patients considered for sample collection.

| **Clinical/Demographic Parameters** | **All patients, n=24** |
|---|---|
| Age(Years) | 65±11.03 |
| Race | |
| Caucasian | 21 |
| African American | 3 |
| Sex | |
| Male | 17 |
| Female | 7 |
| % tumor [enrich] | 98.4±6.24 |
| % nuclei [enrich] | 77.2±11.23 |

#### Reference

1. Robertson AG, Kim J, Al-Ahmadie H, Bellmunt J, Guo G, Cherniack AD, et al. Comprehensive Molecular Characterization of Muscle-Invasive Bladder Cancer. Cell. 2017;171(3):540-56 e25.
